# Causal modeling of gene effects from regulators to programs to traits: integration of genetic associations and Perturb-seq

**DOI:** 10.1101/2025.01.22.634424

**Authors:** Mineto Ota, Jeffrey P. Spence, Tony Zeng, Emma Dann, Alexander Marson, Jonathan K. Pritchard

## Abstract

Genetic association studies provide a unique tool for identifying causal links from genes to human traits and diseases. However, it is challenging to determine the biological mechanisms underlying most associations, and we lack genome-scale approaches for inferring causal mechanistic pathways from genes to cellular functions to traits. Here we propose new approaches to bridge this gap by combining quantitative estimates of gene-trait relationships from loss-of-function burden tests with gene-regulatory connections inferred from Perturb-seq experiments in relevant cell types. By combining these two forms of data, we aim to build causal graphs in which the directional associations of genes with a trait can be explained by their regulatory effects on biological programs or direct effects on the trait. As a proof-of-concept, we constructed a causal graph of the gene regulatory hierarchy that jointly controls three partially co-regulated blood traits. We propose that perturbation studies in trait-relevant cell types, coupled with gene-level effect sizes for traits, can bridge the gap between genetics and biology.

## 1 Introduction

Genome-wide association studies (GWAS) and rare variant burden tests have identified tens of thousands of reproducible associations for a wide range of human traits and diseases. These signals have identified many genes that can serve as therapeutic targets [1–3]; driven discoveries of new molecular mechanisms [4, 5], critical cell types [6] and physiological pathways of disease risks or traits [7–9]; and enabled genetic risk prediction for complex diseases [10].

But despite these successes, it remains difficult to interpret the vast majority of associations. Aside from coarse-grained analyses such as identifying trait-relevant cell types and enriched gene sets, we lack genome scale approaches for interpreting the molecular pathways and mechanisms through which hundreds, if not thousands, of genes impact a given phenotype.

One challenge for interpreting genetic associations is the observation that many hits act indirectly, via trans-regulation of other genes [11–16]. This observation is formalized in the omnigenic model [17, 18] which proposes that, for any given trait, only a subset of genes, referred to as core genes, are located within key molecular pathways that act directly on the trait of interest. Meanwhile, many more genes impact the trait indirectly, by regulating core genes through links in gene regulatory networks. In this model, we can interpret the effect size of a variant in terms of all paths through the network by which it affects core genes.

The central role of trans-regulation underlying many GWAS hits implies that fully understanding the genetic basis of complex traits requires tools to measure how genetic effects flow through networks. But, until recently, we have had very limited information about gene regulatory networks in any human cell type, with the main information coming from observational data such as trans-eQTL and co-expression mapping [11, 13, 19]. However both approaches have important limitations including low power [18, 20] and confounding effects of cell type composition [11] in the case of trans-eQTLs, and ambiguous causality in co-expression analysis [21, 22].

Advances in genome editing and single cell RNA sequencing, including Perturb-seq, now provide new opportunities to measure causal gene-regulatory connections at genome-scale [23–26]. In Perturb-seq experiments, a pool of cells is transduced with a library of guide RNAs, each of which causes knockdown (or other perturbation) of a single gene. After allowing the cells time to equilibrate, single cell sequencing is used to determine which genes were knocked down in each cell and measure the cell’s transcriptome. Critically, Perturb-seq enables measurement of the trans-regulatory effects of each gene in a controlled experimental setting at genome-wide scale. Recent work has shown that such approaches are a promising tool for interpreting GWAS data, finding that GWAS hits are often enriched in specific transcriptional programs identified by CRISPR perturbations of subset of genes [27–31].

Yet, so far it has been challenging to go beyond identifying enriched programs to inferring genome-scale causal cascades of biological information. In this paper, we aimed to develop a new systematic approach to this problem. We demonstrate how, by combining loss-of-function (LoF) burden results with Perturb-seq, we can infer an internally coherent graph linking genes to functional programs to traits, and derive biological insight into the key genes and pathways that control these traits (Figure 1A). The resulting graph helps us to understand not only the trait-relevant pathways but also the functions of genes and programs within the graph, to explain *why* those genes are associated with the traits. Based on our results here, we expect that forthcoming efforts to generate perturbation data in a wide variety of cell types will provide a critical interpretative framework for human genetics.

**Figure 1:**
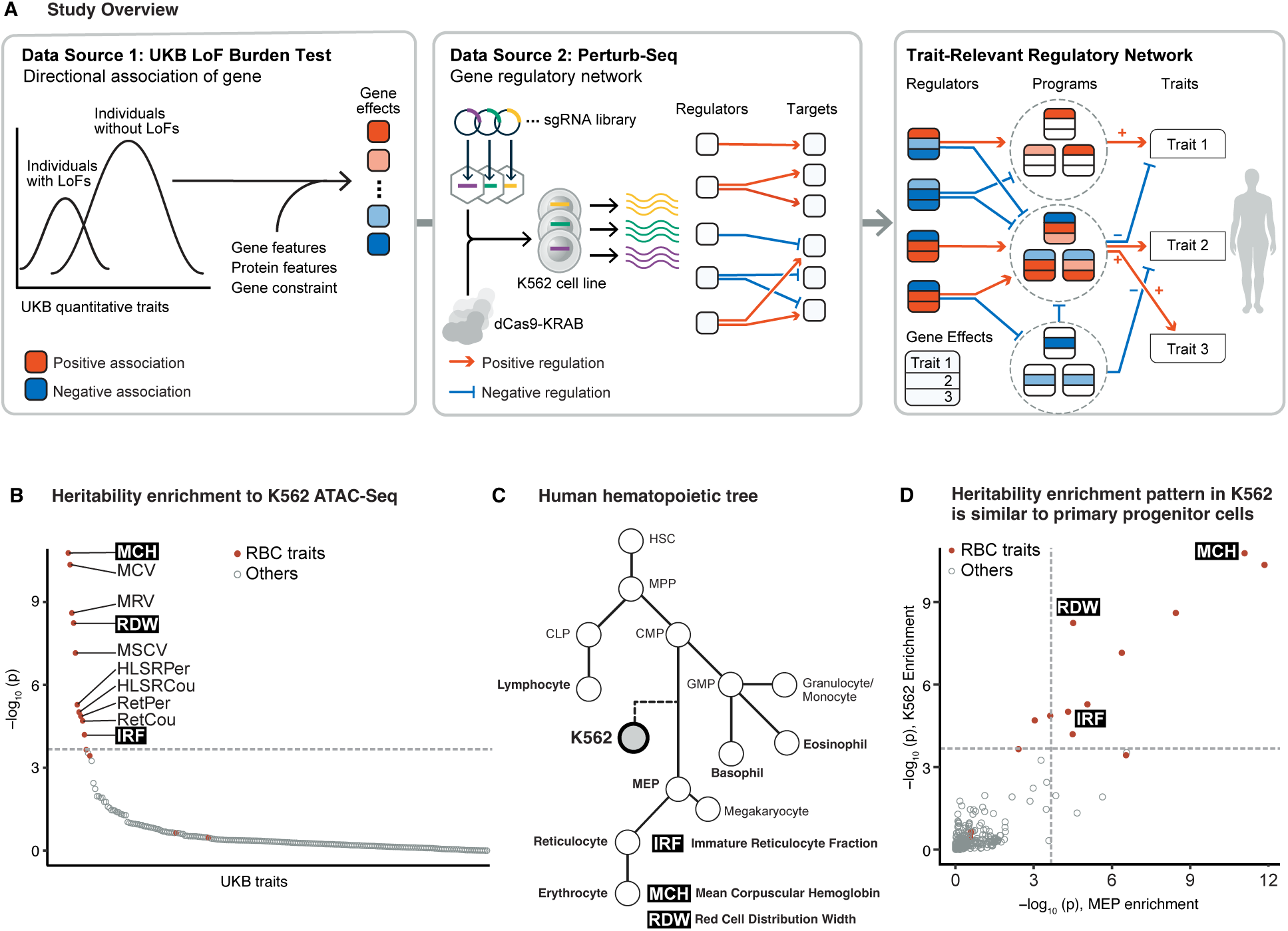
Study overview and selection of model traits **A)** Overview of the study. Square nodes represent genes. Colored arrows between genes represent regulatory effects. Arrows from genes to traits represent associations. **B)** Heritability enrichment of UKB traits to open chromatin regions in K562. Traits are ordered based on the p-value of enrichment from S-LDSC. **C)** Schematic of the human hematopoietic tree. HSC, hematopoietic stem cells. MPP, multipotent progenitors. CMP, common myeloid progenitors. MEP, megakaryocyte/erythroid progenitors. CLP, common lymphoid progenitors. GMP, granulocyte-monocyte progenitors. Traits of interest are annotated near their relevant cell types. **D)** Comparison of heritability enrichment to UKB traits, between MEP and K562 open chromatin regions.

## 2 Results

### Selection of model traits

To integrate genetic association data with Perturb-seq, our first step was to evaluate whether there are any traits with high quality genetic data where the most relevant cell type(s) can be well-modeled by existing Perturb-seq data. At the time of writing, the only published genome-wide Perturb-seq data set was collected in a leukemia cell line, K562 [32]. In that experiment, every expressed gene was knocked down using CRISPR interference, one gene per cell, prior to single cell RNA-sequencing.

To determine which traits could reasonably be modeled in terms of the gene regulatory networks of K562 cells, we compiled published GWAS and LoF burden test data for a wide range of traits measured in the UK Biobank (UKB) [33, 34]. Of these, we selected 468 traits with SNP heritability > 0.04 for further consideration, and performed stratified LD score regression (S-LDSC) [6] across all 468 traits. We observed that open chromatin regions in K562 exhibited significant heritability enrichment exclusively for traits related to morphology or quantity of erythroid lineage cells (Figure 1B). This result is intuitive, as the K562 cell line was derived from erythroleukemia cells, which are a neoplastic form of erythroid progenitors (Figure 1C), and K562 cells retain multipotency and can differentiate into erythroid cells [35].

We also performed S-LDSC across the same set of traits for various primary cell types, and found a very similar enrichment for erythroid traits in megakaryocyte-erythroid progenitor cells (MEP), which are the natural progenitor cells for erythrocytes (Figure S1A, Figure 1C,D). The open chromatin regions in MEP were also more similar to those in K562 cells than other cell types (Figure S1B). These results support the notion that K562 cells share similar chromatin features with primary progenitor cells and could serve as a cellular model for studying the gene regulatory network associated with erythroid traits.

Among the enriched traits, we selected three traits that are relatively independent, with pairwise genetic correlations ranging from -0.39 to 0.15, for detailed analysis (Figure S1C). We focus primarily on *mean corpuscular hemoglobin (MCH)*, which measures the mean amount of hemoglobin per erythrocyte; but we also analyze *red cell distribution width (RDW)*—the standard deviation of the size of erythrocytes per individual—and the *immature reticulocyte fraction (IRF)*.

In addition to serving as model traits for using Perturb-seq to interpret association signals, these three traits are also clinically relevant. They are differentially affected by various causes of anemia [36], and high RDW reflects poor quality control in erythrocyte differentiation, and is associated with mortality in human cohorts [37].

### Pathway enrichments for blood trait associations

Before attempting to build causal models for these traits, we first explored the genetic associations for MCH, RDW, and IRF with standard approaches (Figure 2, S2). GWAS of MCH in the UKB identified 634 independent genome-wide significant signals. Many of the lead hits fall into a few significantly enriched pathways, including heme metabolism, hematopoiesis, and cell cycle (Figure 2A,C). These enriched pathways are crucially involved in the maturation of erythrocytes. For example, tight control of cell cycle is important at several steps in erythropoiesis [38–41].

**Figure 2:**
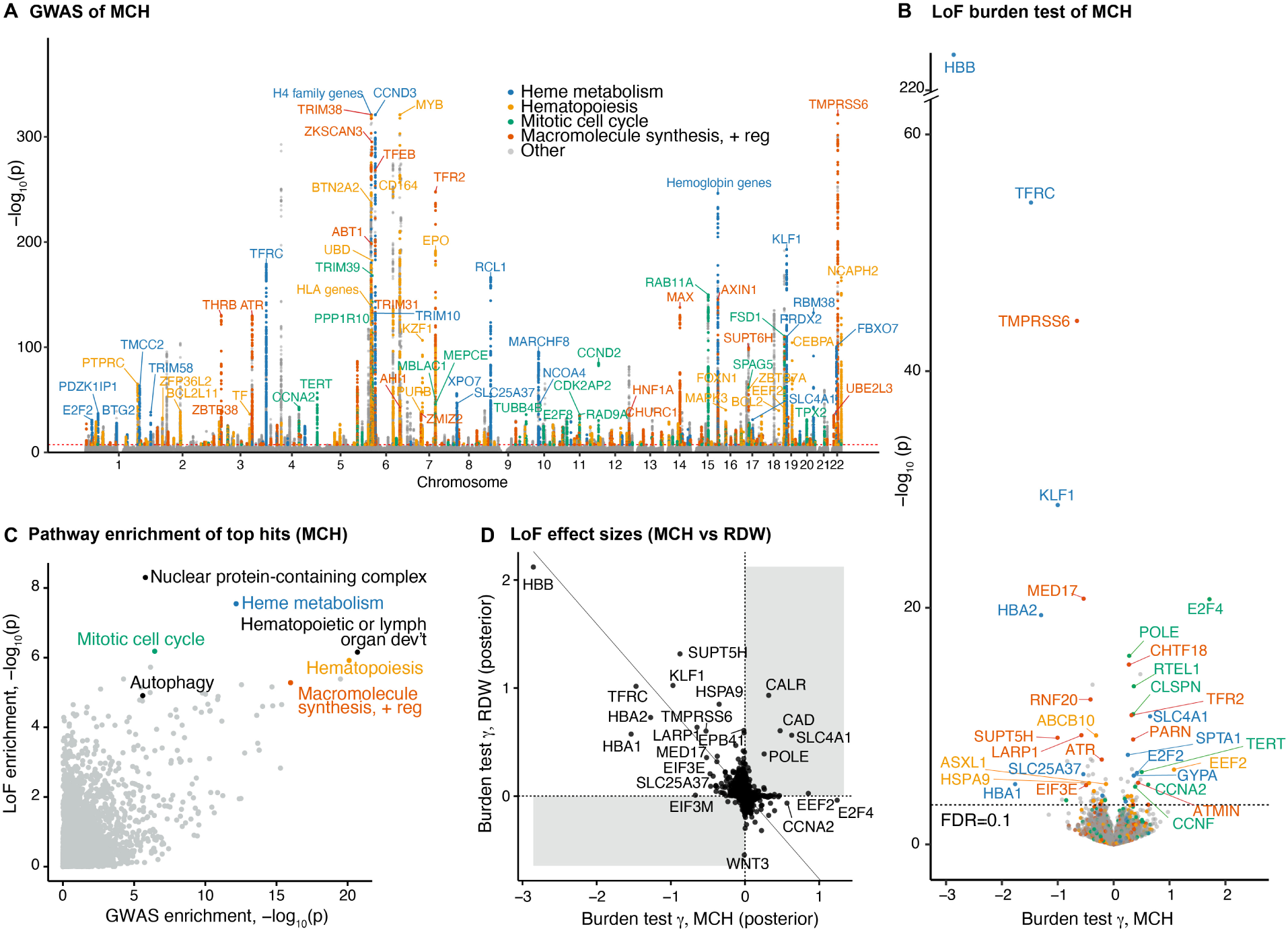
Pathway enrichments for blood trait associations. *Genetic associations identified from UKB GWAS for MCH. Variants located within a 100 kbp window centered on the transcription start site of the genes in the gene set are colored. “Macromolecule synthesis + reg” refers to the “positive regulation of macromolecule biosynthetic process”. **B)** Gene associations with MCH from UKB LoF burden tests. Colors indicate the same gene sets as A). Labeled genes have FDR < 0.01 and belong to the gene sets. **C)** Pathway enrichment of GWAS and LoF burden test top genes. For GWAS, the closest genes from the independent top variants were used. For the LoF burden test, genes were ranked by the absolute posterior effect size from GeneBayes, and the same number of genes as in GWAS was used. **D)** Comparison of LoF burden test effect sizes after GeneBayes between MCH and RDW. The solid line corresponds to the first principal component*.

In addition to GWAS, UKB has also released whole-exome sequencing data for more than 450,000 participants [42]. Here, we focused on the phenotypic effects of LoFs, which are variants such as frameshift and premature stop mutations that are predicted to cause complete loss of function of a gene. The average effects of LoFs on each gene are estimated using a burden test; intuitively, this test estimates the effect of heterozygous loss of gene function on a trait.

Previously-reported burden test statistics for LoF variants [34] identified 90 genes associated with MCH at an FDR=0.1 (Figure 2B). Although the rankings of top hits differ between GWAS and LoF burden tests (Figure S3A), the lead hits from GWAS and LoF are generally enriched in the same pathways (Figure 2C). This is consistent with the expectation that common and rare variants associated with a trait act through similar biological pathways, but frequently prioritize different genes [43, 44].

As one might expect, LoF variants in the genes that code for components of adult human hemoglobin, *HBB*, *HBA1* and *HBA2*, all show strong negative effects on MCH (Figure 2B). Clinically, these mutations cause alpha or beta thalassemia, where a decrease in MCH is characteristic. This highlights a key feature of burden tests: in addition to significance testing, they also provide a quantitative, directional estimate of LoF effects, referred to here as *γ*.

The directions of associations in the burden tests also help us interpret the pleiotropic effects of genes. When looking at genes associated with MCH and RDW, which have a negative genetic correlation in GWAS (Figure S1C), the LoF effects for most genes were also associated in opposite directions (r = −0.53, Figure 2D). However, a handful of genes had strong *same*-direction effects on both traits (Figure 2D). For instance, *CAD* encodes a multifunctional enzyme of which biallelic mutations cause megaloblastic anemia [45], while heterozygous LoFs increase both MCH and RDW (Figure 2D). One goal of building a causal mechanistic graph for these traits will be to explain these seemingly discordant associations.

For many genes, the LoF *γ*s have large standard errors, due to the low frequency of LoF variants [43]. To improve estimation of the *γ*s, we applied an empirical Bayes framework called GeneBayes that we developed recently [46]. Our approach incorporated prior information about gene expression, protein structure, and gene constraint to share information across functionally similar genes (Methods). We find that the GeneBayes estimates of *γ* are far more reproducible than naive estimates in the independent All of Us cohort [47] (Figure S3B). Furthermore, we observed greater enrichment of genes associated with traits in functional pathways even though we did not directly use that information (Figure S3C,D). These improvements are important for making full use of the beneficial features of LoF burden tests while reducing unwanted noise. Therefore, we use the GeneBayes posterior mean effect sizes in Figure 2C, D and for the remainder of the paper.

### Regulatory effects in Perturb-seq explain genetic association signals

Next, we investigated whether Perturb-seq from K562 could allow us to interpret genetic associations in the context of the gene regulatory network. Perturb-seq estimates the effect of knocking down a gene *x* on the expression of another gene *y*, which we denote *β_x_*_→y_ (Methods). *β_x_*_→y_ represents the total effect of *x* on *y*, including both direct and indirect pathways through the gene regulatory network. Previous studies using perturbations to interpret GWAS have identified enrichment of hits in co-regulated gene sets, sometimes referred to as “programs” [27–31], but have had limited success at identifying GWAS enrichment among program regulators (Supplementary note).

As an initial proof-of-concept, we focused on the genes encoding constituents of adult hemoglobin. We focused on the gene *HBA1*, which is the only one abundantly expressed in K562 cells, and which has one of the largest LoF effect sizes for MCH (*γ_HBA1_* = −1.5). We reasoned that if K562 Perturb-seq is relevant for interpreting MCH, then genes that regulate *HBA1* should also be associated with MCH. Moreover, we should be able to predict the direction of effect on MCH from the Perturb-seq data: positive regulators of *HBA1* should, themselves, have promoting effects on MCH, and vice versa for negative regulators. (Note that we refer to genes with negative *β* or negative *γ* from knockdown or LoF, respectively, as *promoting* and color them red; positive *β* and *γ* are considered *repressing* and colored blue.)

As predicted, we found that across all 9, 498 perturbed genes, the LoF effect of a gene *x* on MCH, denoted *γ_x_*, is significantly positively correlated with that gene’s knock-down effects on *HBA1* expression, *β_x_*_→*HBA1*_ (*p* = 3 × 10^−7^, Figure 3A). Notably, among the perturbed genes, of the top ten genes ranked by LoF effects on MCH, seven had nominally significant Perturb-seq effects on *HBA1*, and for all seven the sign of the Perturb-seq *β* matched what we predicted from *γ*.

**Figure 3:**
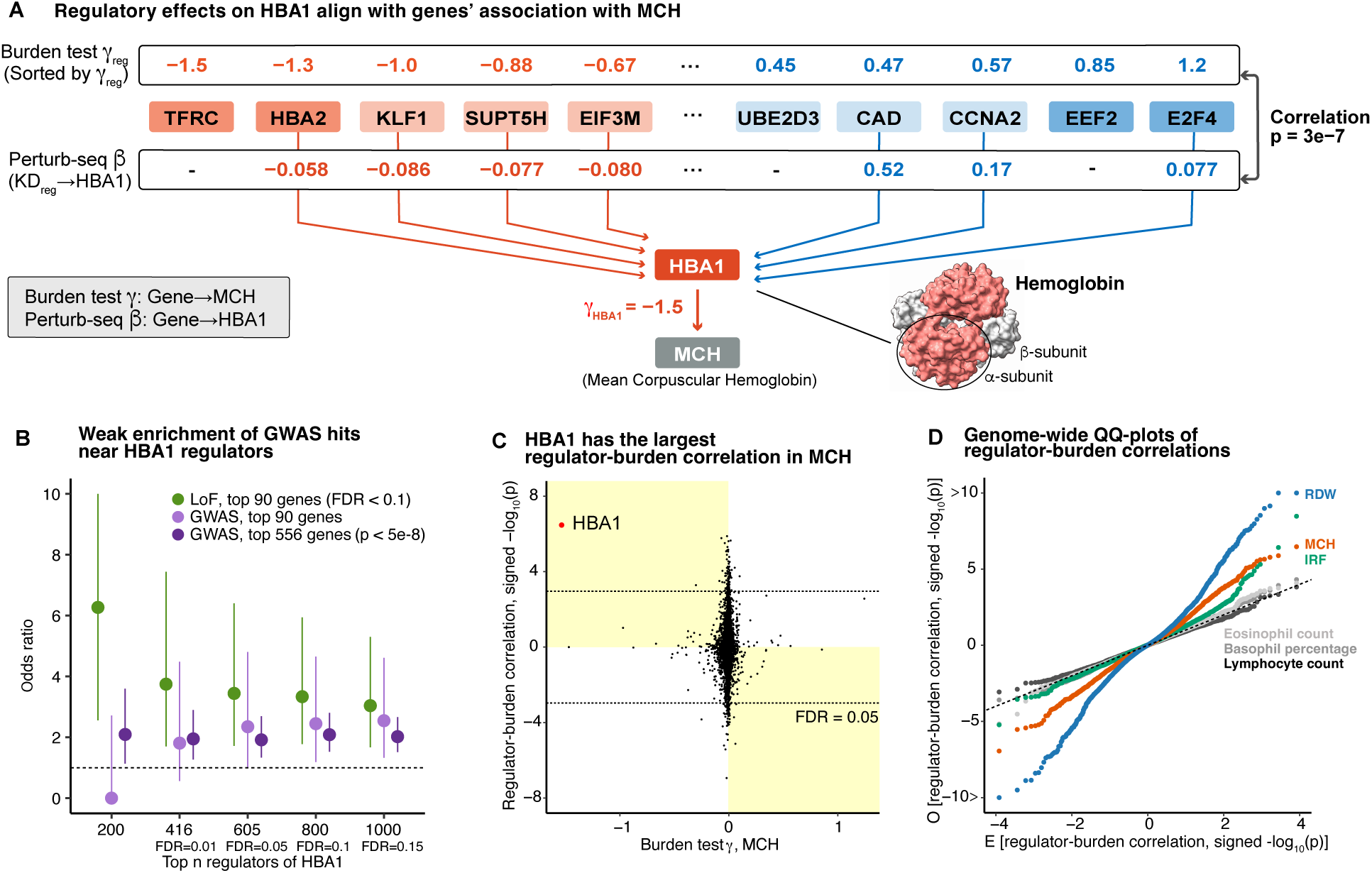
Regulatory effects in Perturb-seq explain genetic association signals. ***A)*** *Gene effects on MCH can be predicted by regulatory effects on HBA1. Genes perturbed in Perturb-seq experiment are ordered by their effect sizes on MCH from LoF burden test. Perturb-seq β refers to log fold change of HBA1 expression after knockdown of the genes. Significant (p < 0.05) regulatory associations in Perturb-seq are connected with arrows. Protein structure of hemoglobin is presented using UCSF ChimeraX* [48] *based on Protein Data Bank entry 1A3N. **B)** Enrichment analysis testing whether the top n HBA1 regulators (ranked by p-values), are enriched at LoF or GWAS top hits. GWAS hits are the closest genes to the independently associated variants (Method). Error bars indicate 95% confidence intervals. **C)** For every expressed gene in K562, regulator-burden correlation is plotted against their γ for MCH. The y-axis shows the -log*_10_*(p) of the regulator-burden correlation, multiplied by the sign of the correlation. Quadrants with yellow background correspond to “concordant” association, where the signs of regulator-burden correlation aligns with the sign expected from the γ of the gene. **D)** Genome-wide QQ-plots for burden-regulator correlations among representative traits. Each dot represents one gene. Traits without significant signals lie along the dotted line. E, expected. O, observed. For other traits, see Figure S4C*.

We also attempted a similar analysis for GWAS hits, testing whether significant GWAS hits were enriched near *HBA1* regulators (Figure 3B). We observed that GWAS hits were enriched (OR = 2.1 for the top 200 regulators), but to a lesser extent than for significant LoF burden test hits (OR=6.3 for the top 200 regulators). This cannot be solely explained by inaccurate gene-linking, as the same set of GWAS hits showed high enrichment for some of the gene sets (Figure 2C, Figure S4A). This suggests a benefit of LoF burden tests over GWAS for identifying the trait-relevant regulatory networks.

We were curious whether similar patterns of correlation between LoF effect and perturb-seq regulatory effects might be found for other genes, or other traits. Consistent with the central role of *HBA1* in determining the MCH phenotype, we found that the correlation of *γ_x_* with *β_x_*_→y_, which we call regulator-burden correlation, was the highest for *y* = *HBA1* among all genes expressed in K562 cells (Figure 3C). As a negative control, we also tested for correlations between regulatory effects on *HBA1* with LoF effects on unrelated traits. As expected, we only detected *HBA1* regulator signals for erythroid traits (Figure S4B). These *HBA1* regulator signals for traits were also detected if we used raw burden effect estimates without applying GeneBayes, but with weaker significance (Figure S3E,F), supporting our approach.

Another key question for Perturb-seq studies is whether regulatory relationships learned in one cell type—K562 in this case—are useful for studying traits that are determined by less-related cell types. To examine this, we computed the regulator-burden correlation for all expressed genes, with LoF *γ*s for a variety of traits. For each trait, we visualized the distribution of regulator-burden correlations in a two-sided QQ-plot.

Starting with our three main erythroid traits, MCH, RDW, and IRF, we see that all three traits show large excesses of both positive and negative correlations compared to the null (x=y line), indicating significant relationships between Perturb-seq and LoF burden tests for many genes. In contrast, there was minimal correlation between regulatory effects and *γ* for other blood traits including lymphocyte and eosinophil counts (Figure 3D, Figure S4C). This suggests that cell types that are not differentiated from MEPs cannot be modeled well using K562 cells (Figure 1C). This observation implies the importance of obtaining Perturb-seq data in trait-relevant cell types.

However, we were surprised to see that some non-erythroid traits, including serum levels of IGF-1 and CRP, as well as BMI, did show highly significant correlations of regulatory effects with *γ* (Figure S4C). The strongest correlations were seen for IGF-1 (insulin-like growth factor 1), which connects the release of growth hormone to cell growth, acting on many cell types and promoting cellular proliferation [49]. Further examination revealed that these signals appear to be driven by genes involved in cellular proliferation including *MKI67*, which is a widely used marker of proliferation. We hypothesize that a cell proliferation network containing *MKI67* may be broadly shared across cell types that regulate IGF-1 and other traits that share this signal (Figure S4D).

Together, these results confirm the relevance of gene-regulatory relationships learned from Perturb-seq for interpreting complex traits. They highlight the role of both cell type-specific pathways—for which the cell type used in Perturb-seq must be closely matched to the trait of interest—and broadly active pathways that may be detectable in many cell types.

### Association of program regulation with blood traits

We next aimed to develop a more comprehensive framework to explain genotype-phenotype associations in terms of the regulatory hierarchy inferred from Perturb-seq data. In principle, one might imagine inferring a complete gene regulatory network from Perturb-seq that contains all causal gene-to-gene edges. However, the inference of accurate genome-scale causal graphs is extremely challenging, if not infeasible, from current Perturb-seq data.

As a more robust alternative, we followed previous work by clustering genes into co-expressed groups, referred to here as programs [28]. To identify programs, we applied consensus nonnegative matrix factorization (cNMF) [50] to the gene expression matrix from Perturb-seq (Figure 4A). This allowed us to quantify the activity of each program in every cell. Similar to [28] we then used the perturbation data to estimate the causal regulatory effects of knockdown of every gene *x* on the activity of each program *P*, denoted *β_x_*_→*P*_.

**Figure 4:**
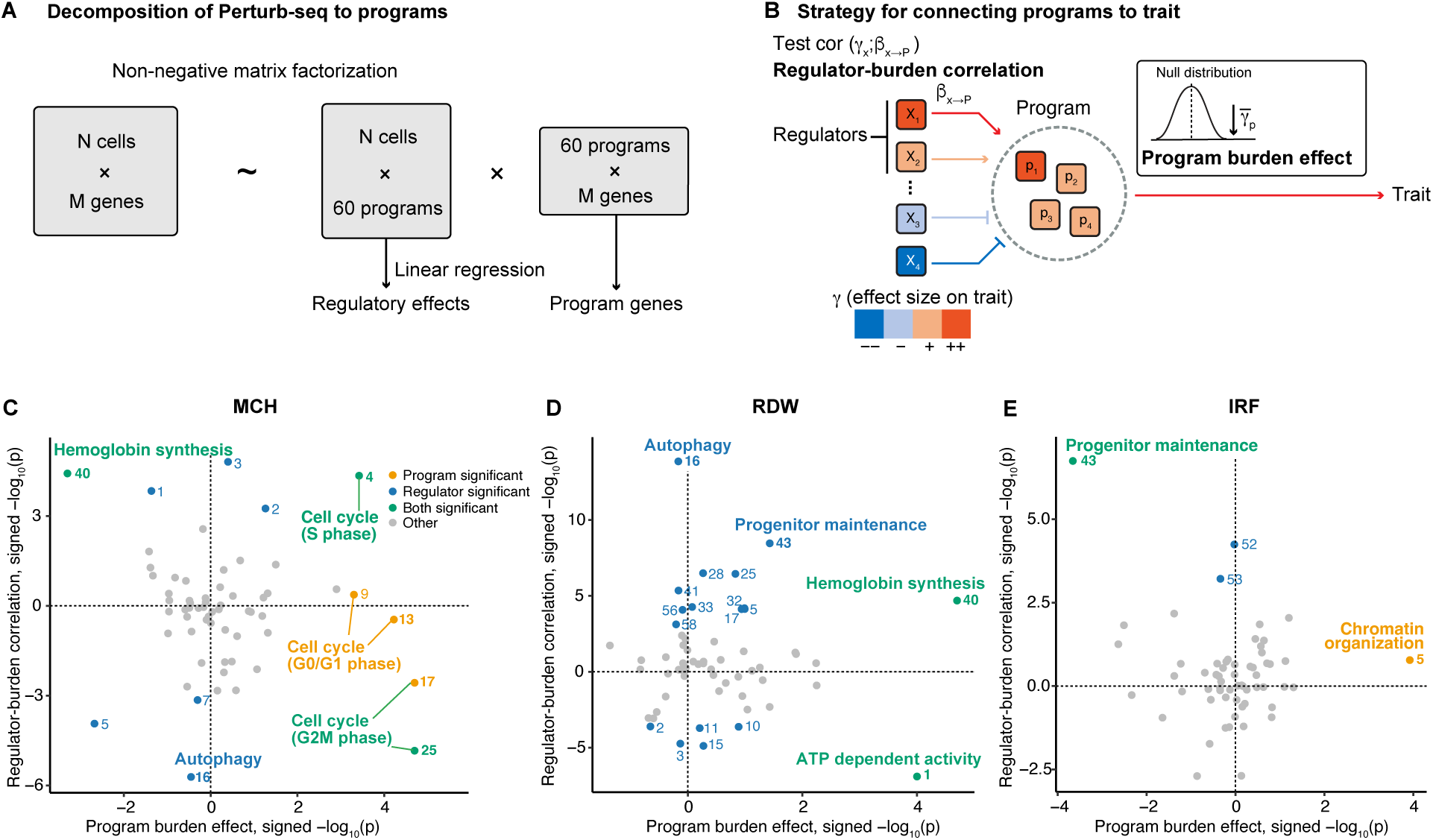
Association of program regulation with blood traits. ***A-B)*** *Overview of our pipeline for the analysis to find the trait-relevant programs. **C-E)** Program burden effects (x-axis) and regulator-burden correlation (y-axis) of 60 programs in three blood traits. Programs with significant associations after Bonferroni correction (P < 0.05/60) are colored. Pathway annotations of representative programs are labeled. For annotations of other programs, see Table S1*.

Based on preliminary analyses, we chose to model the data using 60 programs (Methods, Figure S5A). We found that the 60 programs successfully captured biological pathways, and all were enriched for at least one Gene Ontology category (Table S1). Using external ENCODE data, we found evidence for coordinated transcriptional control of most programs: for 55 of the 60 programs at least one TF showed significant binding site enrichment near program genes and knockdown of that TF significantly changed program expression (Figure S5B, Table S1) [51].

We next quantified the average effects of programs, and their regulators, on traits (Figure 4B). To measure program effects, we note that in non-negative matrix factorization, the gene loadings on each program are non-negative by definition. Thus, a natural measure of a program’s effect on a trait is simply to compute the average LoF effects (*γ*s) of highly-loaded genes as a measure of the effect of that program on the trait. We refer to this as the *program burden effect*. A positive program burden effect is interpreted to mean that the program has a repressing function on the trait; a negative value implies it is promoting. Significance was determined by permutations (Methods).

To measure the effects of regulators of program *P* on each trait, we need to account for the fact that distinct regulators can have either positive or negative effects on *P*. Thus, for each program *P* we computed the correlation across regulators, *x*, of *β_x_*_→*P*_ with *γ_x_*. We refer to this measure as the *regulator-burden correlation*; this measure is analogous to the measure of regulatory effects used for single genes above. A positive regulator-burden correlation is interpreted to mean that up-regulation of program *P* promotes the trait; a negative value suggests that up-regulation of *P* has a repressing effect on the trait.

The program effects on each trait are shown in Figures 4C-E. For MCH, the hemoglobin synthesis program genes and their regulators were both significantly enriched, consistent with our single-gene analysis of *HBA1*. Additionally, five programs associated with the cell cycle were all enriched in the program burden effect axis. This mirrors the enrichment of this pathway from the over-representation analysis of GWAS and LoF top hits (Figure 2C), but here we can confirm the enrichment of both regulators and program genes for these programs (Figure 4C). We will discuss the directional concordance of the programs and regulators in the next section.

For RDW, the program reflecting ATP-dependent activity was highlighted from both program and regulator axes (Figure 4D). This is consistent with previous observations that mitochondrial dysfunction results in sideroblastic anemia, characterized by high RDW [52]. For IRF, the program representing the maintenance of the erythroid progenitor population was enriched for both program and regulator axes (Figure 4E). This program showed the enrichment of binding sites for TFs that are important for the maintenance of stem cell and progenitor populations, including TAL1, NFIC, MAX and MNT [53–55] (Figure S5B).

Overall, the Perturb-seq data efficiently captured biological pathways and their regulators, and comparison with gene associations enable us to identify the pathways relevant to each trait.

### The association of traits, programs, and their regulators reveals complex biology

While the significant programs in Figure 4 provide insight into biological controls of these three traits, they also revealed puzzling inconsistencies. Some programs, including hemoglobin synthesis for MCH, show consistent directional effects for program genes and program regulators, but for other programs the directions of effects initially appeared to be inconsistent (Figure 5A). Examination of these programs revealed important principles about the regulatory architecture of programs, and design considerations for building regulatory models of complex traits.

**Figure 5:**
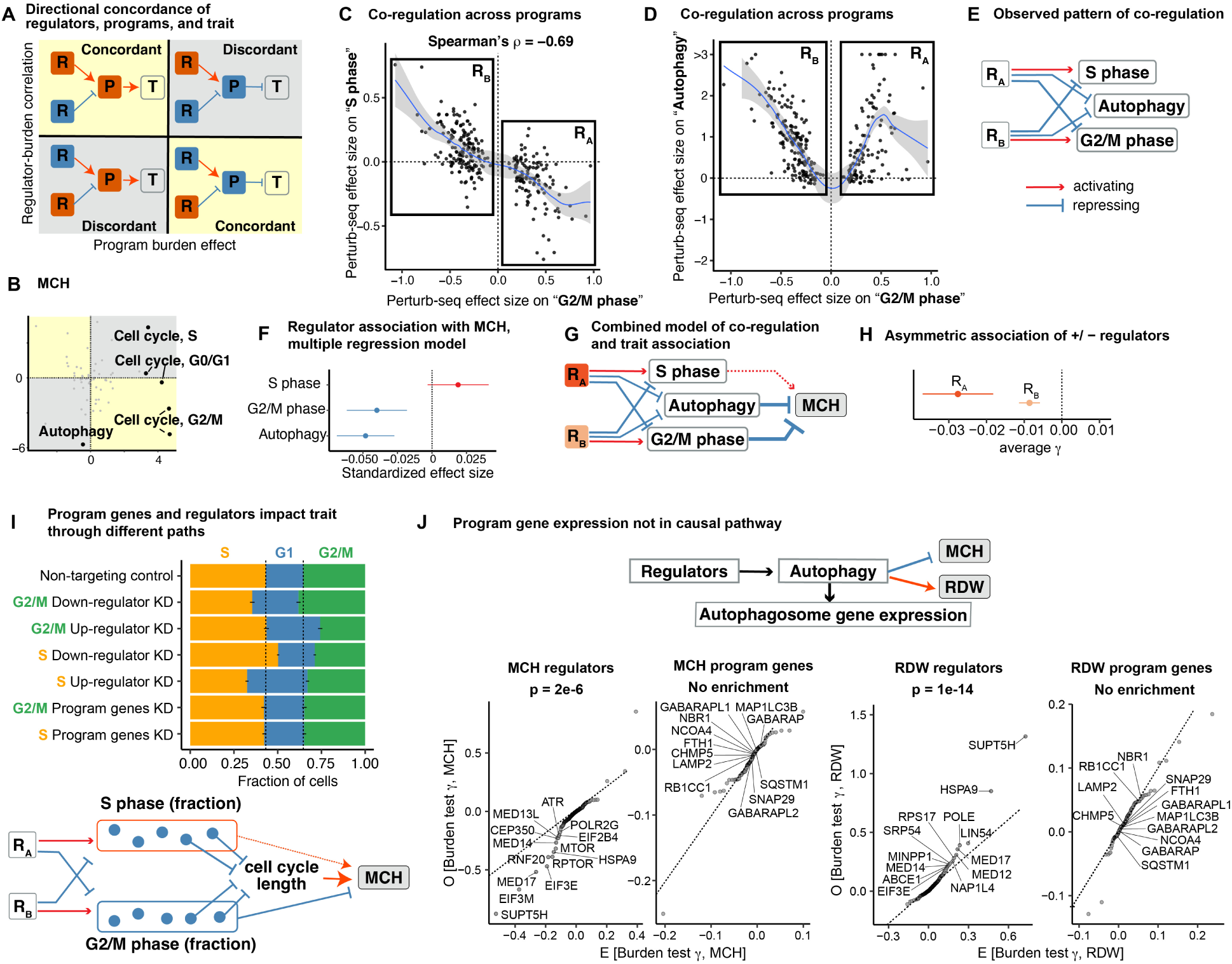
The association of traits with programs and their regulators reveals complex biology. ***A)*** *Schematics for the concordant and discordant patterns between program burden effects and regulator-burden correlation. R, regulators. P, programs. T, trait. **B)** Program association plot for MCH, as shown in* Figure 4B*, highlighting programs discussed here. **C-D)** Co-regulation patterns between programs. Each dot represents a gene that has significant regulatory effects on the G2/M phase program. **E)** The summary of signs of regulatory effects on the programs. **F)** Correlation of regulatory effects on three programs with MCH γ in the multiple regression model. Error bars indicate 95% CI. **G)** Model that combines the co-regulation pattern and trait association of programs. **H)** Observed average γ for MCH. Error bars indicate standard errors. **I)** The fraction of cells in different cell cycles in the groups of cells with perturbations (top) and the model for explaining the cell cycle program association with MCH (bottom). Error bar indicates standard error estimated from Jackknife resampling. **J)** Distribution of burden test effect sizes for MCH (left) and RDW (right). Plots show negative regulators of the autophagy program (FDR<0.05) and the top 100 genes for autophagy program by loading weights (see Figure S6E for positive regulators)*.

### Cross-talk between regulators

The first principle is revealed by three programs with strong effects on MCH: the S and G2/M phase cell cycle programs, and the autophagy program. For G2/M phase, the program and regulator effects have directionally concordant effects on MCH, but for S phase, the program genes and their regulators imply effects with opposite directions. And for autophagy, only the regulators—but not the program genes—show a signal (Figure 5B).

One piece of this puzzle is explained by considering patterns of co-regulation across the three programs: i) regulators of the S phase and G2/M phase programs are shared but affect the programs in opposite directions (Figure 5C); and ii) most G2/M and S phase regulators also affect autophagy, but the knockdown effect on autophagy is almost always positive (Figure 5D, Figure S6A). These relationships are intuitive: S and G2/M are mutually exclusive components of the cell cycle; meanwhile, autophagy is suppressed during mitosis, and cell cycle regulators are known to play a key role in that suppression [56].

To describe these patterns in a simple way, we defined two sets of regulator genes, denoted *R_A_* and *R_B_*, according to their effects on G2/M. The relationships between *R_A_* and *R_B_* and the three downstream programs are summarized in Figure 5E.

To determine how the regulators of these three programs affect MCH, we fit their effects jointly in a multiple regression model. This modeling tests the effects of regulator genes, as mediated independently through each program. This analysis showed that G2/M and autophagy regulators both have independent repressive effects on MCH (Figure 5F). S phase regulators generally promoted MCH, however their effects on the S phase program are smaller in magnitude and colinear with their effects on G2/M, and not significant in multiple regression. The opposite co-regulation of S and G2/M phase programs explains the opposite correlation of these regulators with *γ* (Figure 5B). A summary of the joint model of regulator effects is shown in Figure 5G.

One prediction of this model is that *R_A_* regulators should have stronger (more negative) genetic effects on MCH *γ*s than *R_B_* regulators. This is because *R_A_* genes have a repressive effect on both G2/M and autophagy, and both programs have repressive effects on MCH; while for *R_B_* the positive regulator effects on G2/M and negative regulator effects on autophagy partially cancel each other’s effects on MCH. Indeed, consistent with this model, we see that both *R_A_* and *R_B_* have significantly negative *γ*s on average, but *R_A_* is much more strongly negative (Figure 5H).

These observations emphasize the need for joint modeling of programs, and show that observed effect sizes of regulators on a trait can be modeled as sums of regulatory effects as mediated through key pathways. A different form of cross-talk between programs involving a negative feedback loop affecting RDW is shown in Figure S6D (see Supplementary note).

### Functional differences between program activity and program genes

However, these observations do not yet explain all the observed patterns. Specifically, why do genes that regulate the S phase program promote MCH, but genes *within* the S phase program repress MCH (Figure 5A,B)? And why is there a strong repressing signal for genes that regulate the autophagy program, but genes within the autophagy program have no average effect?

To better understand the functional impact of regulators and program genes on cell cycle, we inferred the cell cycle phase for single cells in the Perturb-seq data set (Figure 5I). When we looked at regulators, these generally had the expected effects: for example, knockdown of negative regulators of S phase increased the number of cells in S phase, and decreased the number of cells in G2/M phase.

However, unexpectedly, knockdown of program genes for both G2/M and S phase had minimal effects on cell cycle proportions. Yet, we did see a strong signal that knockdown of program genes reduced cell growth in a separate K562 growth screen (Figure S6B,C) [57]. Together, these results suggest a functional distinction between genes that *regulate* S phase, and genes *within* the S phase program: knockdown of regulators changes the duration of S phase in the predicted directions, while knockdown of program genes causes functional defects in mitosis without necessarily changing the time in S phase.

To connect these observations to MCH, we note that MCH is strongly positively correlated with cell size [36]. Cell size, in turn, is strongly determined by cell cycle efficiency including correct pausing at key maturation steps and overall cell cycle speed [38–40, 58]. During erythropoiesis, a disrupted (slower) cell cycle leads to a decrease in the number and an increase in the size of erythrocytes [41], leading to higher MCH [36].

In summary, we propose that the regulators and program genes are associated with MCH through different effects on the cell cycle. Regulators change the fractions of cells in each phase and thus affect progression through the maturation process. An increase in G2/M or decrease in S phase fractions results in smaller MCH. Meanwhile, knockdown of G2/M and S program genes both cause defects in cell cycle-related processes, leading to slower cell cycle, larger cells and higher MCH (Figure 5I). Crucially, here we see that the effects of knockdown of genes inside the cell cycle programs differ from knockdown of genes that regulate cell cycle program activity.

A second example of functional differences between program activity and program genes is seen for autophagy. For both MCH and RDW, the regulators of the autophagy program showed higher regulator-burden correlations than any other program (Figure 4C, D). The program genes were highly enriched in the autophagosome pathway (Gene Ontology *P* = 4 × 10^−9^), and yet these genes did not show a significant program burden effect for either MCH or RDW (Figure 5J).

The autophagosome is a cellular structure that delivers organelles to lysosomes during autophagy, which is an essential step in erythrocyte maturation [59]. Development of the autophagosome is precisely regulated by many conserved molecules [59], but the autophagosome itself does not have a degradative function, and its components, represented by the GABARAP gene family (Figure 5J), appear to have some functional redundancy for its formation [60]. Unlike for cell cycle program genes, knockdown of autophagosome program genes does not have significant effects on cellular growth (Figure S6C). In summary, here we see that program gene expression acts as a “proxy” of the pathway activity, but is not in the causal path from regulators to trait.

Together, these observations highlight key modeling considerations. First, gene associations with traits are often sums of multiple regulatory effects, so we need to jointly model regulatory effects on multiple programs. Second, program genes and their regulators can have distinct relationships with a trait, so it is necessary to model them separately to understand trait-relevant regulation.

### Unified graphs linking genes to programs to traits

We next aimed to build regulatory maps that link genes, programs, and traits into coherent, unified models. Our goals in doing so are twofold: 1) we want to understand, in compact form, the main molecular processes that control a set of traits; and 2) we want to interpret, and even predict, the directions of effects of important trait-associated genes.

For each trait, we selected the top-ranked programs by program burden effects and, separately, in a joint regression model, the top ranked programs by regulator-burden correlations (Figure S7,S8, Methods). Based on our analysis above, we allowed programs and program regulators to have independent effects in the model. After model selection, this procedure resulted in a graph that, for MCH, included 5 programs and 3 sets of program-regulators, as well as the inferred direction of effect of each program and regulator set on MCH (Figure S9).

A simplified representation of the MCH graph is depicted in Figure 6A, showing hemoglobin synthesis, cell cycle, and autophagy as critical controls of MCH. The direction of the genetic association of top genes on MCH was generally consistent with this model (43 out of 59 predicted correctly). Importantly, the overall prediction accuracy was significantly higher than expected under a null model, using both leave-one-out cross validation (p = 5 × 10^−5^) and permutation analyses for which we repeated the entire inference procedure (p < 5 × 10^−5^; Methods, Figure S10A, S10B). This approach allows us to connect the gene-level top hits, identified solely from genetic association studies, to their functions in the pathway regulatory map.

**Figure 6:**
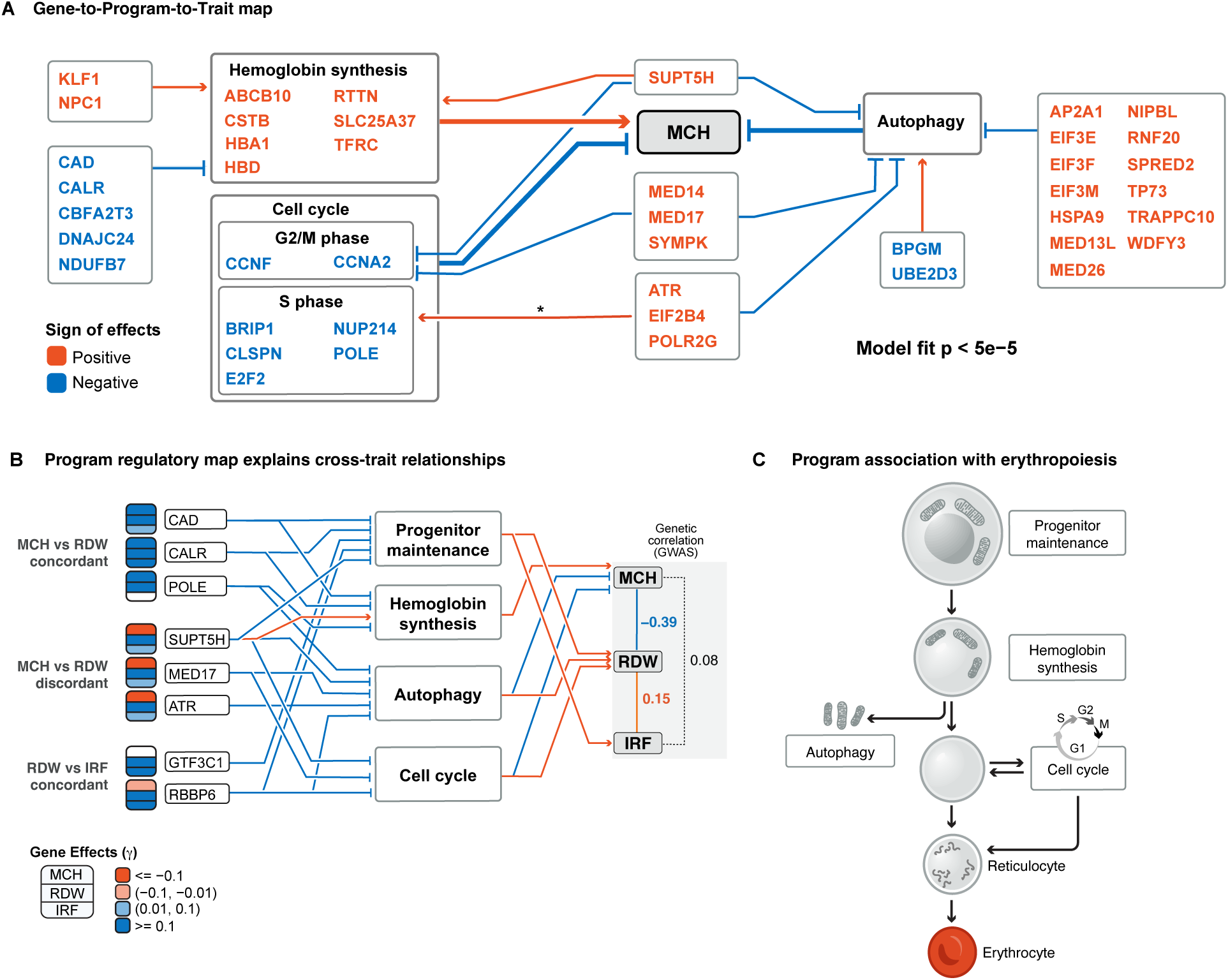
Association map of genes to programs to traits. ***A)*** *Regulatory map of MCH. Programs were selected by genome-wide association patterns of regulators or program genes with the trait (Methods). Top hits for MCH (*|*γ*|*>0.1) whose effect directions were concordant with the model are placed onto the map. Color of the genes indicate the direction of effects on the trait (sign*(*γ*)*); red, increase MCH with up-regulation of the gene. Arrow with * was not selected in the initial program selection process. **B)** Sharing of regulatory networks across traits. Here, the arrows from gene to programs indicate the regulatory directions. Programs were selected if their regulators were found to be associated with at least one trait in the gene-to-program-to-trait map. Arrows from programs to traits were determined based on joint regression model (Figure S11C). Regulatory directions on cell cycle pertain to G2/M phase. POLE is also a member of S phase cell cycle program. **C)** Programs identified in our model are associated with biological processes that are essential for erythrocyte maturation*.

Examining the graph, we were intrigued that *SUPT5H*, which is involved in transcriptional elongation [39], has regulatory effects on all three programs. Perturb-seq shows that *SUPT5H* activates hemoglobin synthesis, and inhibits autophagy and the G2/M phase cell cycle, all of which result in increased MCH (Figure 6A). Thus, our model predicts that *SUPT5H* is a master regulator for MCH, exerting same-direction effects via three different pathways. Indeed, the effect sizes of *SUPT5H* LoFs on MCH are among the largest of all genes (Figure 2D), and LoFs in this gene can cause a thalassemia phenotype [61]. Thus, this map can help us to interpret why genes are associated with a trait.

In addition to MCH, we also inferred gene-to-pathway-to-trait maps for RDW and IRF, revealing both shared and independent pathways of regulation across the three traits (Figure S10C-H). There were four programs whose regulators were significantly and independently associated with at least one trait (Figure 6B): progenitor maintenance, hemoglobin synthesis, autophagy, and cell cycle. Previous studies of hematopoiesis confirm that all four pathways regulate essential aspects of erythrocyte maturation (Figure 6C) [39, 41, 62, 63].

The multi-trait regulator graph (Figure 6B) helps us to interpret the concordance and discordance of genetic associations across the traits. Genome-wide, MCH and RDW are negatively correlated in both GWAS data (*r_g_* = −0.39) and at significant burden loci (Figure 2D). We can now interpret these observations as likely driven by opposite direction effects of both autophagy and cell cycle on these two traits. Conversely, RDW and IRF are positively correlated (*r_g_* = 0.15; Figure S11A), at least in part because both traits are positively regulated by progenitor maintenance.

We can also use the graph to understand how individual genes affect the different traits. For example, 16 genes in the graph have strong opposite-direction effects on MCH and RDW; our model correctly predicts opposite signs for 14 out of the 16, including *SUPT5H*, *MED17*, and *ATR* (Figure 6B, Figure S11B). For instance, *MED17* inhibits both G2/M phase cell cycle and autophagy; both effects increase MCH and reduce RDW, with the result that *MED17* increases MCH and reduces RDW.

In contrast, three genes in the graph differ from the genome-wide pattern, showing large same-direction effects on MCH and RDW. Our model correctly predicts two of these, and is suggestive for the third (*POLE*; Supplementary Note). Specifically, *CAD* and *CALR* both have repressive effects on RDW and MCH. Figure 6B suggests why: unlike most genes that affect both RDW and MCH through shared pathways, these genes affect the two traits via independent pathways, namely progenitor maintenance and hemoglobin synthesis. Both genes inhibit both pathways, but regulation of progenitor maintenance affects RDW and not MCH, while hemoglobin synthesis affects MCH but not RDW.

While the trait graphs were inferred from LoF burden signals, we tested whether we could have identified similar patterns using enrichment of GWAS hits. As we saw for hemoglobin (Figures 3B, S4A), there was generally strong enrichment of GWAS hits near program genes, but enrichment of regulators was modest (Figure S12A-C). This affirms the value of LoF signals for this type of analysis.

In summary, by using Perturb-seq and LoF burden tests, we can construct detailed graphs that allow us to interpret and predict the effects of genes based on how they affect the expression or function of the programs.

## 3 Discussion

Genetic associations serve a unique role in studies of human biology, as they can establish causal links from variants or genes to human traits and diseases. Yet, some 20 years after the first GWAS, we still lack genome-scale approaches for inferring interpretable, quantitative models of the biological pathways that connect genes to cellular functions to traits. Here, we build on previous work in this area [27–31] to develop the first approach to infer unified graphs linking directional effects of genes on traits via pathways of regulation and cellular functions. While our work focuses on blood traits that underlie anemia and related diseases, we anticipate that the principles learned here can be broadly applicable.

One essential feature of this paper is that we build graphs using quantitative gene effects estimated from LoF burden tests instead of unsigned enrichment of GWAS hits. We envisage LoFs and GWAS hits as reflecting the same underlying biological pathways [43, 44], but our results are both more significant, and more interpretable, when using LoFs. Unlike GWAS hits, LoF effect sizes are inherently directional, they are automatically linked to the correct genes, and their magnitudes are comparable across genes. Moreover, compared to common variants with tiny effects, LoFs are likely more functionally similar to CRISPR knockdowns, given the widespread nonlinear and even non-monotonic relationships between gene expression and phenotypes [64, 65].

Use of signed effects to build a causal graph also provides further advantages over simply testing for program enrichment. First, we can perform more stringent model testing—for example, asking if we can predict the directions of genetic effects in cross-validation. Second, the signed effect sizes provide far more biological insight than simple enrichment—for example, showing specifically that autophagy is a negative regulator of MCH rather than showing merely that autophagy is enriched, and explaining why particular genes have either concordant or discordant relationships between MCH and RDW. Third, the use of directional effects in the model was essential for revealing a striking discordance between regulators and programs, as we observed for S phase genes, and for explaining why two sets of regulators of cell cycle (denoted *R_A_* and *R_B_*) have 3-fold different effect sizes on MCH.

While the model presented here is relatively simple, there will surely be value in future models that add complexity. Future versions could allow for more complex representations of gene regulatory networks, more explicit modeling of regulatory cross-talk between programs, and heterogeneity of gene functions within programs. Many traits are controlled by multiple cell types, and one can envision models in which genetic effects on traits are controlled by a superposition of effects across multiple cell type-specific networks.

One unexpected result from our model was the finding that the effects of program regulators on a trait may be strongly discordant from the effect of program genes on the same trait, as we observed for S phase and autophagy. We hypothesize that some programs reflect downstream transcriptional consequences of cell biological processes, and that the genes within a program do not always lie on the causal pathway between the program-regulators and the trait. In such cases, the identification of genes in the program can provide useful clues about biological mechanism but the effects of program genes may differ dramatically from the effects of their regulators. Moreover, it is likely that some critical processes may not be detected or may not be interpretable from RNA readouts. Thus it will be helpful in future analyses to augment Perturb-seq experiments with other types of cell phenotyping such as functional tests, protein measurements, or cell painting [66–69].

Lastly, one critical challenge for using Perturb-seq to interpret association studies is how closely we need to match the cells used for Perturb-seq to the cells that determine trait variation [28]. Recent work suggests that gene regulatory relationships are often shared between closely-related cell types, but generally not shared between more distant cell types [70, 71]. Consistent with this, our results show that K562 serves as a suitable, though imperfect, model for erythrocyte development – but also that K562 is not suitable for modeling traits related to other blood cell lineages (Figure 3D). We hypothesize that in general Perturb-seq data will need to be closely matched to the trait-relevant cell types, but the matching does not need to be perfect.

While our proof-of-principle here uses experimental data from K562 cells to model erythrocyte traits, we expect that the next generation of perturbation studies in cells, organoids, and tissues [69, 72] will provide a critical interpretative framework for human genetics.

## 4 Methods

### Datasets

*GWAS data*. We downloaded the publicly available GWAS summary statistics and SNP heritability estimates for traits in the UKB from Ben Neale’s lab (see URL section below). We focused on traits with SNP heritability estimates exceeding 0.04. For binary traits we used heritability estimates on the liability scale.

*LoF data*. We used LoF burden test summary statistics from the UKB with 454,787 participants, as previously reported [34]. Specifically, we utilized the gene-level aggregated effect estimates from predicted LoF variants with a minor allele frequency of <0.01%. Data were downloaded from GWAS Catalog [73].

*Perturb-seq data*. We utilized the genome-wide Perturb-seq dataset in K562 reported by Replogle *et al*. [32]. In this dataset, all expressed genes (N = 9,867) were targeted by a multiplexed CRISPRi sgRNA library in K562 cells engineered to express dCas9-KRAB. Single-cell RNA-seq was performed to read out the sgRNAs together with the transcriptome. Only cells with a single genetic perturbation were used for the analysis, amounting to a median of 166 cells per gene perturbation and 11, 499 UMIs per cell. We downloaded the raw count data that the authors uploaded to figshare (see URLs section).

*ChIP-seq data*. We utilized ChIP-seq data in K562 for annotating gene programs. We downloaded 830 transcription factor ChIP-seq narrow peak files from the ENCODE project website [51] (see URLs section). All coordinates were mapped to hg19 with LiftOver [74].

### LD score regression

To identify traits whose heritability is enriched in open chromatin regions in K562, we used stratified LD score regression [6]. All GWAS data were preprocessed with the “munge_sumstats.py” script provided by the developers (URLs). Variants in the HLA region were excluded from the analysis. The ATAC-seq narrow peak bed file in K562 was downloaded from ENCODE [51] (GSE170378_ENCFF590CPE_replicated_peaks_GRCh38.bed.gz) and the coordinates were mapped to hg19 using LiftOver [74]. Furthermore, we used narrow ATAC-seq peaks from 18 hematopoietic progenitor / precursor / differentiated cell populations previously reported [75].

LD scores were calculated for each annotation using the 1000G Phase 3 European population reference [76]. The heritability enrichment of each annotation for a given trait was computed by adding the annotation to the baseline LD score model (v1.1) and regressing against trait chisquared statistics for HapMap3 SNPs. These analyses used v1.0.1 of the stratified LD score regression package (URLs).

Further, we tested the genetic correlation between specific trait pairs using European LD scores with the LD score regression package (v1.0.1).

### Estimation of gene effect sizes with GeneBayes

#### Method overview

LoF burden tests are not well-powered, especially for shorter or selectively constrained genes, as the likelihood of having LoF variants in these genes is low. We previously developed GeneBayes [46], an empirical Bayes framework aimed at addressing a similar challenge—the precise estimation of selective constraint on genes, which can be particularly challenging for short genes. Within GeneBayes, we use gene features in a machine learning-based empirical Bayes framework to improve the accuracy of constraint estimates. Diverse gene features, such as gene expression patterns and protein structure embeddings, can enhance the accuracy of these estimates. GeneBayes is a highly adaptable framework, easily extendable to various applications, as outlined in the original manuscript [46]. In this instance, we utilized it to derive more precise effect size estimates for LoF burden tests.

To minimize overfitting when applying GeneBayes to LoF burden test estimates, we first performed feature selection using the BoostRFE function (Boost Recursive Feature Elimination) from the shap-hypetune package (URLs) to fit XGBoost [77] models on the sign and magnitude of *γ*^, the estimated effect size from LoF burden test summary statistics. We used the predicted sign and magnitude as the features for GeneBayes, which we found to perform better than using the selected features directly; this may be due to differences in training dynamics between XGBoost and the gradient-boosted trees fit using GeneBayes.

Subsequently, we implemented the GeneBayes framework as previously described. Specifically, GeneBayes involves two steps: firstly, learning a prior for the effect size of each gene through the utilization of gradient-boosted trees, as implemented in NGBoost [78], and secondly, estimating gene-level posterior estimates of the effect sizes using a Bayesian framework. In our application of GeneBayes, we parameterize the prior as follows:

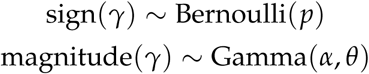

The parameter *p* is the probability that *γ* is positive or negative, and *α*, *θ* are the shape and scale parameters of the Gamma distribution respectively. We learn the parameters of the prior using the following likelihood:

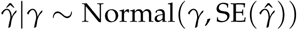

The summary statistics *γ*^ and SE(*γ*^) are the estimated effect size and its standard error from the LoF burden tests respectively.

#### Gene features

We compiled the following types of gene features from several sources: selective constraint of genes (*s_het_*) [46], gene expression across cell types, protein embeddings, and gene embeddings. Gene expression across 79 single cell types was downloaded from the Human Protein Atlas [79] (see URLs). Protein embeddings were adopted from embeddings learned by an autoencoder (ProtT5) trained on protein sequences [80]. Gene embeddings were derived from GeneFormer, a pretrained deep learning model for single-cell transcriptomes [81]. Specifically, we used the Cell×Gene Discover census (see URLs), and we extracted 1,000 cells per each of the following cell types: ’erythroid progenitor cell’, ’monocyte’, ’erythrocyte’, ’fibroblast’, ’T cell’, ’neutrophil’, ’B cell’, ’hematopoietic stem cell’ and computed the average embeddings of each gene for the cellular classifier using the EmbExtractor module (URLs).

Finally, we used the posterior mean of the LoF burden test effect size as a point estimate for the following analyses.

#### Traits

As applying GeneBayes to all UKB traits is computationally intensive, we applied this to a subset of traits including all the blood cell associated traits, blood biomarkers and some of anthropometric traits. A list of traits included in our analyses is in Table S2.

#### Evaluation in the independent cohort

The All of Us cohort has conducted whole genome sequencing and reported the LoF burden test statistics for some of the traits [47]. At the time of writing, the LoF burden test Z-score for “red cell distribution width,” which is equivalent to RDW in UKB, was reported with 114,402 cases (UKB data-field 30070, AoU ID 3019897). We utilized this to evaluate the sign concordance. Specifically, we ranked the genes based on burden test absolute effect sizes in UKB, with or without applying GeneBayes, and tested what fraction of the top N ranked genes had the same sign of associations (sign of Z-score) in All of Us (Figure S3B).

### Pathway enrichment analysis of GWAS and LoF top hits

#### Clumping of GWAS top variants

To identify independently associated GWAS variants, we used Plink (v.1.90b5.3) [82] with the –clump flag, a p-value threshold of 5 ×10^−8^, an LD threshold of r^2^ = 0.01, and a physical distance threshold of 10 Mb. Additionally, we merged SNPs located within 100 kbp of each other and selected the SNP with the minimum p-value across all merged lead SNPs to avoid the false inclusion of genes that have neighbor genes with extremely large effects. This resulted in 634 independent variants associated with MCH. For each independent variant, we annotated the nearest proteincoding gene. To accomplish this, we used the bedtools (v2.30.0) [83] closest module to identify genes that overlap with the variant or have their transcription start site or transcription end site closest to the variant. Finally, we obtained a list of 556 genes possibly associated with GWAS signals for MCH.

#### Pathway enrichment analysis

We aimed to compare the pathways enriched in GWAS and LoF top hits for MCH. As pathways, we utilized all ontology terms in Gene Ontology (GO) [84] with a minimum of 20 genes and a maximum of 2,000 genes, as well as MsigDB hallmark genesets [85] that include the heme synthesis pathway. We utilized enrichGO and enricher functions in clusterProfiler [86] package in R for the analysis.

Among the enriched pathways, genes in “positive regulation of macromolecule biosynthetic process” pathway overlaps significantly with those in the “autophagy” pathway (*p* = 2 × 10^−8^), and thus its enrichment may reflect the relevance of autophagy pathway.

### Comparison of gene regulatory effects with genetic association

#### Estimation of gene-regulatory effects from Perturb-seq

Here, we aimed to estimate gene-to-gene regulatory effects from Perturb-seq. We can assess the total effects of gene knockdown on gene expression by comparing perturbed and non-perturbed cells. After filtering out cells with fewer than 500 genes expressed and genes expressed in fewer than 500 cells, we compared the cells with perturbation of every gene versus the cells with nontargeting control gRNAs. Log-normalized counts of cells were used as input to the limma-trend pipeline [87], while accounting for GEM group (batch effect), number of genes expressed, and the percentage of mitochondrial gene expression as covariates.

#### Correlation of regulatory effects with genetic associations

We utilized the log-Fold Change (logFC) of gene expression in perturbed cells compared to non-targeting cells as a point estimate of the perturbation effect on gene expression. We started from a simple model where the effect size of a peripheral gene *x* is determined by its regulatory effects on a limited set of core genes. In cases where there is a single or a limited number of core genes *y*, the regulatory effect size of the peripheral gene on the core genes should correlate with the peripheral gene’s effect size on the trait. We have previously observed a striking correlation between LoF burden test effect sizes and *s_het_* on average across traits [43]. To avoid the confounding effects of selective constraint, we included *s_het_* as a covariate in our linear regression model:

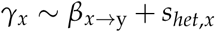

where *β_x_*_→y_ corresponds to the regulatory effect of gene *x* on gene *y*. Gene *y* itself was excluded from the vector of gene *x*. For every expressed gene *y*, we evaluated the significance of the coefficient for the first term. In some of the plots, the significance level was multiplied by the sign of the coefficient.

#### Enrichment analysis of GWAS/LoF top hits to HBA1 regulators

For the evaluation of the GWAS top hits’ enrichment related to *HBA1* regulators ((Figure 3B), we used the list of 556 closest genes to the independent GWAS hits defined above. We ranked the genes based on the p-values of their regulatory effects on *HBA1* expression. We evaluated the enrichment using a two-sided Fisher’s exact test, using all the genes perturbed in the Perturb-seq as a background. Additionally, for comparison, we evaluated the enrichment of 90 significant genes in the LoF burden test (FDR < 0.1) and the genes closest to the top 90 independent GWAS hits.

### Comparison of programs and their regulatory effects with genetic associations

#### Identification of gene programs with cNMF

From a single-cell gene expression matrix, we can identify the co-regulated set of genes. Intuitively, such a set of genes can correspond to genes that determine cellular identity or specific cellular processes, which we call programs. To identify gene programs and their activity in each cell, we applied the consensus non-negative matrix factorization (cNMF) [50] method to the singlecell gene expression matrix from Perturb-seq. With cNMF, the gene expression data matrix can be modeled as the product of two matrices: one corresponding to the contribution of each gene to each program, and the second corresponding to the activity of each program in each cell. In cNMF, a meta-analysis of multiple iterations of NMF is performed to obtain a “consensus” result. In cNMF, the number of programs (*K*) is a key model hyperparameter to tune. To determine it, we tested different values of *K* (30, 60, 90, 120) and decided to proceed with *K* = 60 based on the error versus stability comparison (Figure S5A), as proposed by the authors. Also, we used density_threshold=0.5 to filter out the outlier programs.

#### Annotation of programs to biological pathways

From the gene-by-program matrix produced by cNMF, we can obtain the non-negative loadings of each gene to the program. We ranked the genes based on the loadings and utilized the top-ranked genes for each program to characterize the biological pathways of the program. We examined the enrichment of the top 300 genes in the GO categories using the enrichGO function in the clusterProfiler [86] package in R.

In addition, we can expect that for some programs, the genes within the same program are coordinately regulated by specific transcription factors. Such transcription factors can be used to characterize the programs. To this end, we utilized the ChIP-seq experiments of transcription factors in K562 from the ENCODE project. To convert the information on binding sites to a genelevel regulation score, we calculated the following score for each transcription factor (*i*) for each protein-coding gene (*j*), as adopted from [88]:

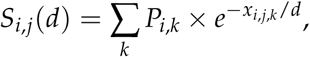

where *P_i_*_,*k*_ denotes the strength of peak *k* for transcription factor *i* (quantified by − log_10_ q-value for each peak, outputted by MACS2), *x_i_*_,*j*,*k*_ denotes the distance from peak *k* to the TSS of gene *j*, and *d* represents the decay distance. The decay distance indicates the effective distance for the transcription factor and can vary depending on the transcription factors. Here, we set the value to either 5 kbp or 50 kbp.

To determine which score is useful for the annotation of programs, we tested the correspondence of the score with differentially expressed genes (DEGs) after KD of the same transcription factor. Specifically, for each transcription factor, we listed positive or negative DEGs after KD in Perturb-seq (FDR < 0.1) and compared the Mann-Whitney *U* test score between DEGs and other expressed genes. As a natural consequence, we can annotate each TF as an activator or inhibitor, according to the direction of effects after KD. We annotated a TF as an activator if the downregulated DEGs after KD had significantly high ChIP-scores (FDR<0.05), and as an inhibitor if the up-regulated DEGs after KD had significantly high ChIP-scores (FDR<0.05). As a result, ChIP-scores for 153 TFs showed significant correspondence with their KD effects and were utilized for the annotation of programs. One of two decay distance parameters was selected for each TF based on the significance in the overlap with DEGs.

For each program, we compared the top 300 loading genes with other expressed genes in K562 with respect to the 153 ChIP-scores using the Mann-Whitney *U* test. This test evaluates the enrichment of binding sites of the TFs to each program genes. Further, we compared the program activity of the TF-KD cells with others to see if the TF has a direct effect on the activity of the program (Figure S5B).

Additionally, we manually confirmed the co-expression of marker genes for predefined cell types or pathways and program activity of cells in Uniform Manifold Approximation and Projection (UMAP) [89] space. Markers for RBCs, myeloid cells, and the integrated stress response pathway were adopted from the original Perturb-seq paper [32]. Markers for erythroid progenitors and megakaryocytes were determined from single-cell gene expression data of bone marrow hematopoietic progenitors [90], where we ranked the genes in each corresponding population based on expression specificity (z-score) compared to other populations, and selected the top 50 genes. S phase and G2/M phase marker gene sets were adopted from [91].

By integrating these sources of information, we annotated each program to a biological pathway (Table S1).

#### Program gene association with traits

Next, we quantified the average effects of program genes on traits, which we call *program burden effect*. Program burden effects are the average *γ* of the genes which are representative of the program, as determined by the loading for the program in cNMF.

Notably, as a feature of cNMF, the loadings of the genes to the programs are always positive. Thus, the sign of the average *γ* provides interpretable directional information about the program association with the trait.

As selective constraints are positively correlated with |*γ*| [43], highly conserved programs, such as those essential for cellular survival, could have larger program burden effects. To avoid confounding, we divided the expressed genes in K562 into 10 bins based on *s_het_*. We then compared the average *γ* of the top loading genes with a 10,000 randomly chosen sets of the same number of genes, while matching for *S_het_* bin. To account for the directional association, we converted the rank of the observed value compared to the random distribution into two-sided p-values, while adding the sign of the average *γ* to calculate the signed association p-values.

The results were generally not affected by the choice of the number of top genes (100, 200, 300). However, for some programs including the hemoglobin synthesis program, where the association with MCH was concentrated on a small number of hemoglobin genes, the association was more pronounced with a smaller number of top genes. Therefore, for Figures 4C-E, we chose 100 for defining the top genes.

#### Regulator association with traits

Next, we aimed to quantify the correlation of regulatory effects of genes on the program with *γ*, which we call *regulator-burden correlation*.

From the cell-by-program matrix produced by cNMF, we can obtain the usage of each program in each cell. To obtain the effect size of each regulator on the program usage, we compared perturbed cells with cells with non-targeting control gRNAs with a linear regression model, while accounting for GEM group (batch effect), number of expressed genes and percentage of mitochondrial gene expression as covariates.

We utilized the point estimate of the effect size of perturbation on program usage as a regulatory effect of a gene. We then calculated the correlation of regulatory effects with trait association signals while accounting for *s_het_*in the same way as the gene-level analysis:

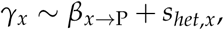

where *β_x_*_→P_ corresponds to the regulatory effect of gene *x* on program *P*.

For visualization of the distribution of burden effects of regulators or program genes (Figure 5J), the expected distribution of burden effect sizes was determined by randomly picking up the same number of genes from non-associated genes 10,000 times and taking their average.

#### Estimation of causal relationships between programs

While examining the co-regulation patterns across programs, we noticed an asymmetric pattern of co-regulation between programs; that is, the regulators of program A also have effects on program B, but the regulators of program B do not have effects on program A (Figure S6D). Such asymmetry can be explained by a causal directional association from one program to the other. Biologically, this one-way association can be interpreted as positive or negative feedback from one program to the other.

A similar observation—that is, the asymmetric correlation of effects from explanatory variables between two traits—was reported in the GWAS literature [92]. For instance, when LDL cholesterol causally affects the risk of coronary artery disease, but not vice versa, the effect sizes for risk variants of LDL cholesterol show a strong correlation between the two traits, while those for risk variants of coronary artery disease do not show such correlation [92].

We adapted the analytic framework for causality from a previous GWAS study [92] to our case. Specifically, for a pair of programs, *p*_1_ and *p*_2_, we identify significant regulators (FDR < 0.05) for each. We then calculate *ρ_p_*_1_, the Spearman’s rank correlation of effect sizes for *p*_1_ and *p*_2_, considering only the regulators of *p*_1_. We can also calculate *ρ_p_*_2_ for the regulators of *p*_2_. Next, we modeled

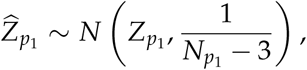

where *Z_p_*_1_ = arctanh(*ρ_p_*_1_) and *N_p_*_1_ corresponds to the number of significant regulators for *p*_1_.

Then we consider four patterns of association, M1: *p*_1_ causally associated with *p*_2_ (*Z_p_*_2_ = 0), M2: *p*_2_ causally associated with *p*_1_ (*Z_p_*_1_ = 0), M3: no relationship between *p*_1_ and *p*_2_ (*Z_p_*_1_ = *Z_p_*_2_ = 0), M4: correlation does not depend on how the regulators were ascertained (*Z_p_*_1_ = *Z_p_*_2_).

We fit each model by maximizing the corresponding approximate likelihood. We then select the model with the smaller Akaike Information Criterion (AIC) from the two causal models (M1, M2) and from the two non-causal models (M3, M4). Finally, we calculate the relative likelihood of the best non-causal model compared to the best causal model.

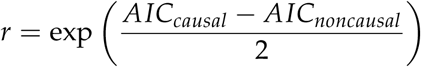

We treat r < 0.01 as a threshold for causally associated programs. In the case of programs associated with RDW (Figure S6D), the causal association from the mitochondrial program to the hemoglobin synthesis program showed *r* = 8.5 × 10^−7^, whereas other pairs of programs had r >

0.05 (also refer to Supplementary Note).

### Validation of multiple program association with the trait

To test whether jointly modeling multiple programs can explain more of the genetic association signals than modeling with a single program, we conducted a cross-validation analysis. We randomly split 80% of the genes into a training set and 20% into a test set, and fitted regression models to explain the gene effects on the trait (*γ*) by gene regulatory effects on the program(s) using the training set. We evaluated the variance of *γ* explained by the model using the test set.

We tested this with the set of multiple programs chosen from the regulator-burden correlations in gene-to-program-to-trait models for MCH and RDW, as well as with the same number of randomly chosen programs, and single program models. The selected multiple program model explained much more variance than any single program model or random combination of programs for MCH and RDW (Figure S7). For IRF, only one program was chosen from the regulator-burden correlation in the gene-to-program-to-trait model, so we did not perform the comparison.

### Construction of Gene-to-Program-to-Trait model

Prevalent co-regulation across programs, as well as feedback, suggested the need to jointly model multiple programs to identify those whose regulation independently explains the trait association signals. Additionally, although program burden effects and regulator-burden correlation sometimes converge on the same program, we have observed cases where either only program content or only regulators are enriched in trait association signals, as well as cases where both program content and regulators are enriched but through different mechanisms. Therefore, we treated program burden effects and regulator-burden correlation separately to identify trait-associated programs included in the model.

#### STEP 1: Selection of programs based on regulator-burden correlations

To select programs whose regulators are enriched for the trait association signals, we conducted a stepwise linear regression analysis using the ‘regsubsets’ function in the ‘leaps’ package [93] in R. In this analysis, we included gene regulatory effects on 60 programs (*β_x_*_→p_), as well as levels of gene constraint (*s_het_*, as defined in [46]) as potential explanatory variables, with *γ_x_*as the dependent variable. We identified the combination of explanatory variables through exhaustive search to determine the best subsets for predicting *γ_x_*in a multiple linear regression model. The number of variables to include in the final model was decided by assessing the variance explained in the model upon changing the number of variables (Figure S13A), along with the significance of the model fit in the subsequent permutation test (Figure S13B). For the MCH model, we opted to include three variables together with *s_het_*: regulators for autophagy, hemoglobin synthesis, and G2/M phase cell cycle programs.

#### STEP 2: Selection of programs based on program burden effects

For selecting programs with enriched contents for the trait association signals, we followed the following process. Firstly, for each program, we calculated program burden effects. That is, we ranked the genes based on their loading and selected the top 200 genes and calculated the average of *γ* of these genes. Then we compared it with randomly selected 10,000 sets of genes expressed in K562 while matching for 10 bins of *s_het_* to calculate two-sided enrichment p-values. Subsequently, we ranked the programs based on these p-values. To determine the number of programs to include in the final model, we varied the number of top programs included and evaluated the model fit in the subsequent permutation test (Figure S13B). Specifically, for the MCH model, 5 programs were selected: the hemoglobin synthesis program and 4 programs associated with different phases of cell cycle. These 5 programs largely corresponded to those that had significant program burden effects after Bonferroni correction in the previous test (Figure 4C).

#### STEP 3: Predicting the signs of associations for the regulators/ program genes in the model

After selecting programs from both regulator and program content associations with the trait, we assigned the predicted signs of effects to each gene in the model. Specifically, for regulators, we considered genes that exhibited significant regulatory effects on the selected programs (FDR < 0.05). In cases where a regulator had regulatory effects on multiple programs, we calculated the gene’s total effects on the model by summing the product of the effect sizes of the selected programs on the trait in the multiple linear regression model (*w_p_*) and the gene effects on the program (*β_x_*_→p_, Figure S8). The sign of this product was utilized as the regulatory direction of the gene to the trait predicted from the model.

For program contents, we assigned the sign of the association of the program (i.e., the sign of the average *γ* of the top loading genes) to the top 200 loading genes. If a gene belongs to both program and regulator genes, although such case was relatively rare, we assigned the sign from the program enrichment test because of the potentially larger effect sizes of program function on the trait (Supplementary note).

#### STEP 4: Assessing the directional concordance of the associations of top hits with the model

To assess how well the predicted model can explain the directional genetic associations, we evaluated it in two ways: leave-one-out cross-validation and permutation testing.

For leave-one-out cross-validation, we left out one gene at a time, selected the programs based on program burden effects and regulator-burden correlation using the other genes, and predicted the sign of the left-out gene as described above. We then assessed the enrichment of correctly predicted genes among the top hits (genes with (|*γ*| > 0.1)), compared to genes with minimal associations (genes with (|*γ*| < 0.01)), using Fisher’s exact test. In this test, the enrichment is influenced by both 1) the enrichment of the top genes among the genes selected in the model (significant regulators or program genes in the model) and 2) the accuracy of the predicted signs among the genes in the model. Our result for the MCH model showed that the top genes were enriched in both 1) selected genes in the model (OR = 1.8), and 2) sign concordance (OR = 1.9), with an overall enrichment of p = 5 × 10^−5^ and OR = 2.2. This result supported the use of Perturb-seq for predicting the directed gene associations.

For the permutation test, we created 20,000 sets of permuted *γ* by permuting gene labels. We then followed the same program selection and sign assignments processes, while fixing the number of selected programs from both program burden effects and regulator-burden correlation. In each permutation, we counted the number of top genes whose sign of association was correctly predicted by the model and evaluated the enrichment over other genes using Fisher’s exact test. Finally, we compared the Fisher’s test p-value of the observed data to those of the permuted sets and calculated the permutation p-value (Figure S10B, D, F). Similar to leave-one-out crossvalidation, we observed that the observed genetic association data had many more concordant genes, along with a higher ratio of concordant to discordant predicted signs compared to the permuted data (Figure S10A, C, E). The permutation test can evaluate the fit of our model to the genetic association signals.

For the permuted dataset, we slightly modified the way for program selection. Here, instead of matching for *s_het_*, we compared the distribution of *γ_x_* between the top loading genes and randomly selected genes expressed in K562 using the Mann-Whitney *U* test to calculate enrichment p-values. Subsequently, we ranked the programs based on these p-values and selected the same number of top programs. This helps to greatly speeds up the process, although the resulting permutation p-value for the model is potentially conservative.

We ran the permutation tests while differing the parameters for the modeling. The model fit to the data was robust to the choice of the number for defining program genes (100, 200, or 300) and to different thresholds for defining high-effect genes (|*γ*|) (Figure S13C). We chose the number of program genes to be 200 and threshold for |*γ*| to be 0.1 based on the highest fit of the model.

#### STEP 5: Drawing gene-to-program-to-trait map

Finally, we aim to draw a map to interpret the functions of the trait-associated genes. Here, we included all the top hits with |*γ*| > 0.1 whose direction of association was concordant with that predicted from the model into the map (Figure 6A). When regulators have concordant regulatory effects on multiple programs, we included all paths in the map.

### URLs

Neale lab UKB data: http://www.nealelab.is/uk-biobank

Replogle et al. Perturb-seq data: https://plus.figshare.com/articles/dataset/_Mapping_in formation-rich_genotype-phenotype_landscapes_with_genome-scale_Perturb-seq_Replogl e_et_al_2022_processed_Perturb-seq_datasets/20029387

LDSC software: https://github.com/bulik/ldsc

ENCODE database: https://www.encodeproject.org/

shap-hypertune package: https://github.com/cerlymarco/shap-hypetune

Gene expression in single cell types: https://www.proteinatlas.org/humanproteome/single+cell+type

CellxGene Discover census: https://chanzuckerberg.github.io/cellxgene-census/

GeneFormer embedding extractor module: https://geneformer.readthedocs.io/en/latest/geneformer.emb_extractor.html

## Acknowledgments

This research has been conducted using the UK Biobank resource under application number 52374. We utilized the All of Us resource under workspace ID aou-rw-c30ba93b. We are grateful to V. G. Sankaran for valuable feedback on an earlier draft of the manuscript, and J. Engreitz, H. Mostafavi, R. Zhu, R. Lopez, N. Milind, C. J. Smith, and members of the Pritchard lab for helpful conversations, and T. Tolpa for help with the figure design. This work was funded by grants R01HG008140, R01HG011432, and U01HG012069. A. M. received funding from the Simons Foundation, Lloyd J. Old STAR Award (Cancer Research Institute), Parker Institute for Cancer Immunotherapy, Innovative Genomics Institute, Larry L. Hillblom Foundation (grant 2020-D-002NET), Northern California JDRF Center of Excellence, the Byers family, K. Jordan and the CRISPR Cures for Cancer Initiative. M. O. is supported by Astellas Foundation for Research on Metabolic Disorder and Chugai Foundation for Innovative Drug Discovery Science. E. D. is supported by an EMBO Postdoctoral Fellowship.

## Supplementary Material

### Supplementary Notes

#### GWAS enrichment to gene regulatory networks from Perturb-seq

Previous studies have investigated the enrichment of GWAS signals to the programs and regulators identified by Perturb-seq [28–30] (and related papers [27, 31]). Consistent with our GWAS analysis here, the previous studies that aimed to link GWAS to Perturb-seq have generally found significant enrichment at the program level, but limited enrichment of regulators. We discuss some of the key papers here:

A pioneering study by Schnitzler *et al.* [28] tested the enrichment of coronary artery disease GWAS variants to programs and regulators identified from Perturb-seq in telomerase-immortalized human aortic endotherial cells, where they perturbed 1,661 genes close to GWAS hits. They applied cNMF to their dataset and identified five out of 50 programs for which the union of program and regulator genes showed significant enrichment to GWAS-linked genes. Their signal was primarily driven by GWAS hits at the program level, and in total only six regulators overlapped GWAS-linked genes for the five enriched programs. Using relationships between regulators and programs inferred from the Perturb-seq data, they proposed a regulatory pathway that controls key programs, but this only contained two GWAS-linked genes. While the authors successfully validated the roles of these key regulators (*CCM2* and *TLNRD1*) in atherosclerosis etiology, it may be difficult to generalize this approach to connect regulatory pathways to traits at genome-scale.

Yao *et al.* [30] performed Perturb-seq in THP1 cells with or without LPS stimulation, where they perturbed 598 genes in a cell-pooled experiment. They identified three modules of regulators that had similar effects on transcriptional programs. One module (genes that suppress the LPS response) was found to be enriched for polygenic signals for 2 out of 64 tested traits (lymphocyte and neutrophil percentages) in an analysis using sc-linker [94]. Positive regulators of the LPS response did not show enrichment to any of the traits. In contrast, they observed many examples of downstream gene signatures (genes whose expression is affected by KD of a focal gene) that were enriched to various traits/ diseases.

Geiger-Schuller *et al.* [29] performed Perturb-seq in mouse primary dendritic cells, targeting 1,130 genes associated with E3 ligases. They tested the enrichment of modules of regulators to polygenic risk of 60 traits/diseases with sc-linker [94] and MAGMA [95], and they reported that one regulator gene module was enriched across all traits, with relatively higher enrichment to immune mediated diseases. In contrast, program genes showed more trait-specific enrichment patterns.

Although these studies used different experimental and analytical protocols, their observations have similarities: despite trait-specific enrichment of co-regulated gene programs, program regulators did not show enrichment to GWAS signals, or any enrichment that *was* found was not trait-specific.

We propose that the difficulties in identifying trait-specific regulators from GWAS enrichment analyses arise from two aspects. First, for GWAS signals, enrichment tests cannot account for the direction of association of genes. As observed in the studies above [29, 30] and in our analysis, genes often have regulatory effects on multiple programs with different directions. Accounting for these directional effects may be important not only for gaining power in the enrichment analysis but also for understanding the trait-specific enrichment. Second, the cis-effect size of common variants on gene dosage is expected to be smaller than that of LoF variants. Furthermore, the trans effects of genes are not linear in relation to gene dosage for many genes, a phenomenon known as buffering [64, 65]. Thus, the ranking of GWAS top hits based on their effect size on the trans program may not align with the ranking of the cis-gene’s trans effect on the program under the KD experiments. Together, the top hits prioritized by GWAS may not overlap well with the top regulators in Perturb-seq, and LoF burden test could be a good alternative for it.

#### Negative-feedback and co-regulation across pathways explain RDW associations

In RDW, the hemoglobin synthesis program exhibited a discordant enrichment pattern in regulator-burden correlation and program burden effects (Figure 4D). Positive program burden effects indicate that the disrupted hemoglobin synthesis process leads to higher RDW. Considering that RDW is defined as the standard deviation of erythroid size, which quantifies the quality of erythrocytes, this observed direction makes biological sense. However, the regulator-burden correlation indicated that up-regulation of hemoglobin synthesis program results in higher RDW. These resulted in seemingly discordant enrichment pattern. How can we interpret this observation?

Here, regulators of the hemoglobin synthesis program also had effects on the ATP-dependent activity program, the other enriched program. However, regulators of the ATP activity program did not have significant effects on the hemoglobin synthesis program, on average (Figure S6D). This pattern suggests a directional relationship from hemoglobin synthesis program activity to ATP activity, corresponding to the biological negative feedback observed between these two pathways [96]. We applied a causal inference method originally used for interpreting GWAS top variant associations [92] to our case (Methods) and indeed observed that the causal model was 10^6^ times more likely than the non-causal models. Thus, the regulatory patterns in the Perturb-seq data suggested a negative feedback from the hemoglobin synthesis pathway to the ATP activity pathway.

In such case, regulators of the hemoglobin synthesis program impact on RDW through their effect on ATP activity. It can result in discordant regulator enrichment for the hemoglobin program in the marginal test.

In addition, regulators of the ATP activity and autophagy programs are highly shared. The multiple regression model suggested that autophagy regulation has the largest effect on RDW among these programs (Figure S6D).

In general, when trait-relevant programs are connected by this kind of feedback loop and/or co-regulation, it can result in seemingly discordant enrichment.

#### Overlap of program genes and regulator genes in the model

Although not very frequent, some of the genes were both top-loading program genes and significant regulators in the selected model for the trait in the gene-to-program-to-trait map (Figure S9 for MCH). Since we cannot quantify the program gene KD effect on program *function*, unlike its expression levels, it is difficult to predict the total effects of the gene on the trait and the signs of associations in this case.

In such cases, we prioritized the signs of program burden effects, based on the assumption that the gene effects on program functions are generally larger than their regulatory effects on other programs.

For example, *POLE* was one of the top-loading program genes for the S phase cell cycle program. At the same time, it was a negative regulator for both the hemoglobin synthesis program and the autophagy program. These programs were all associated with MCH.

As the sign of association of the S phase program with MCH, predicted from program burden effects, was repressing, the predicted direction of POLE for MCH as an S phase program gene was “repressing”.

On the contrary, the regulator-burden correlation indicated that the quantity of the hemoglobin synthesis program was positively associated with MCH, while the autophagy program was negatively associated. Thus, the regulatory effects of POLE on these two programs canceled each other out, leading to our predicted signs of the effects of POLE on MCH as a regulator being weakly “activating”.

In our model, we predicted that its effect on MCH was “repressing” as we prioritize the signs from program burden effects. In this case, it was concordant with the observed *γ* of this gene on MCH.

In contrast, RDW was only associated with POLE via its regulatory effects on the autophagy program. Also in this case, the sign of association was predicted consistently from the model.

Thus, in this case, the predicted directions of *POLE* associations with the traits were correct for both traits. However, this assumption may not always hold true. For example, a gene co-regulated with other functional program genes may not have direct effects on program functions by itself.

#### Association of *SLC4A1* with MCH and RDW

LoF of *SLC4A1* had the strong effects on MCH and RDW in the same direction (Figure 2D), in contrast to the negative genetic correlation of these traits. *SLC4A1* encodes a cell surface protein expressed on erythrocytes. LoF of one copy of this gene is a known cause of hereditary spherocytosis [97]. Spherocytosis leads to the secondary hemolysis, which can increase both RDW and MCH [97]. In this case, this gene affects these traits not through the regulation of other genes. At the same time, a quantitative change in this gene may not be linearly associated with the trait—for instance, a 20% decrease in gene dosage may not result in hemolysis. The function of this type of gene on the trait cannot be investigated solely through Perturb-seq; experimental and clinical knowledge is needed to complement it.

## Supplementary Tables

Table S1: Annotation of cNMF programs to biological pathways.

Table S2: List of control traits.

## Supplementary Figures

**Figure S1:**
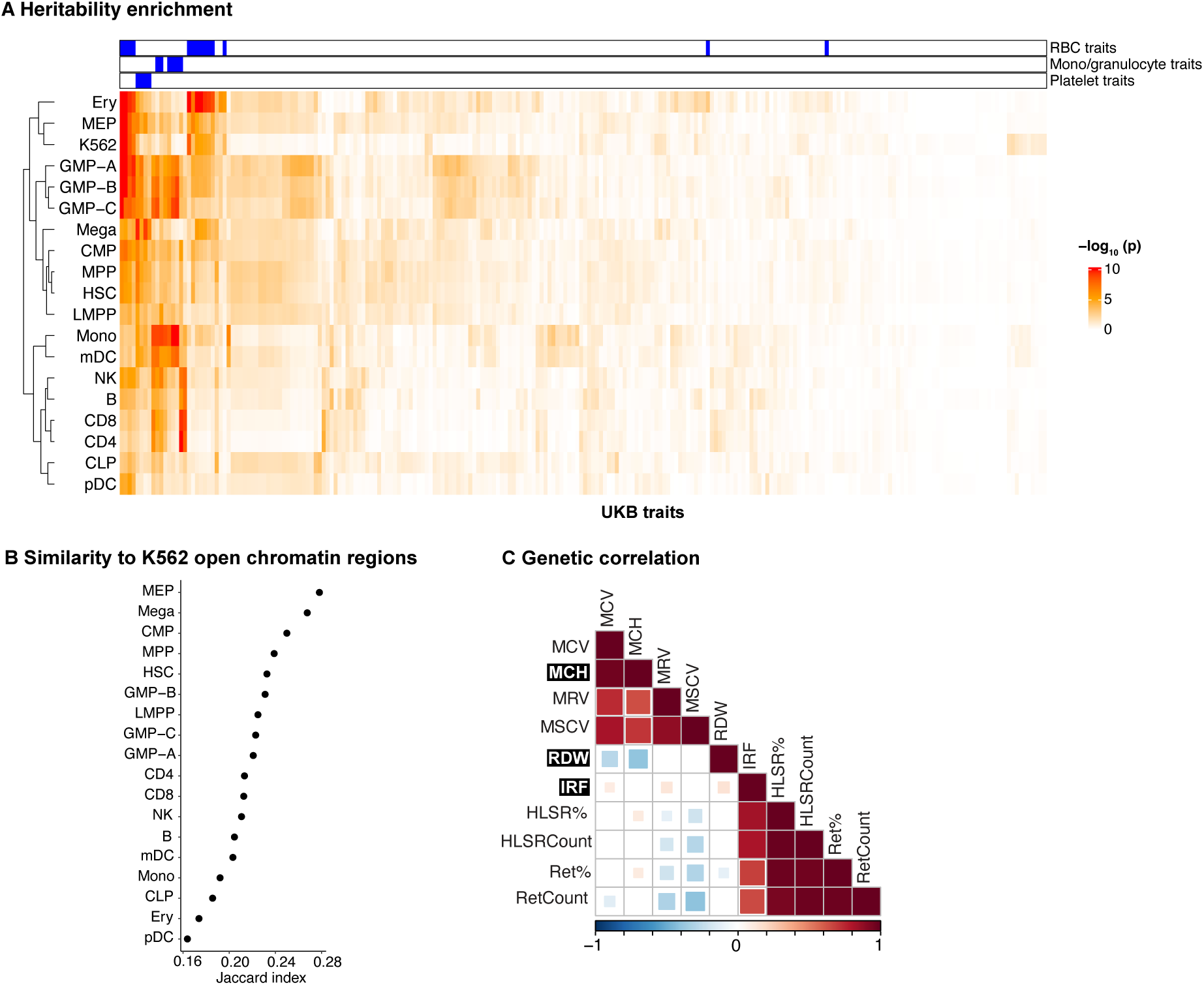
Analysis of the heritability of multiple traits from GWAS, related to. **Figure 1. A)** Heritability enrichment of UKB traits to 18 primary hematopoietic cell types [75] and K562. Heritability enrichment was estimated with S-LDSC by adding each annotation to the baseline model. Traits associated with the morphology or quantity of RBC, monocyte/granulocyte or platelet are labeled on top. Both cell types (rows) and traits (columns) are hierarchically clustered based on their patterns of enrichment. K562 showed the closest similarity to MEP. **B)** Similarity of open chromatin regions of primary cell types to K562. Plotted are Jaccard index, which captures the proportion of open chromatin regions that are shared with K562. **C)** Genetic correlation across traits which were enriched to K562 in S-LDSC analysis.

**Figure S2:**
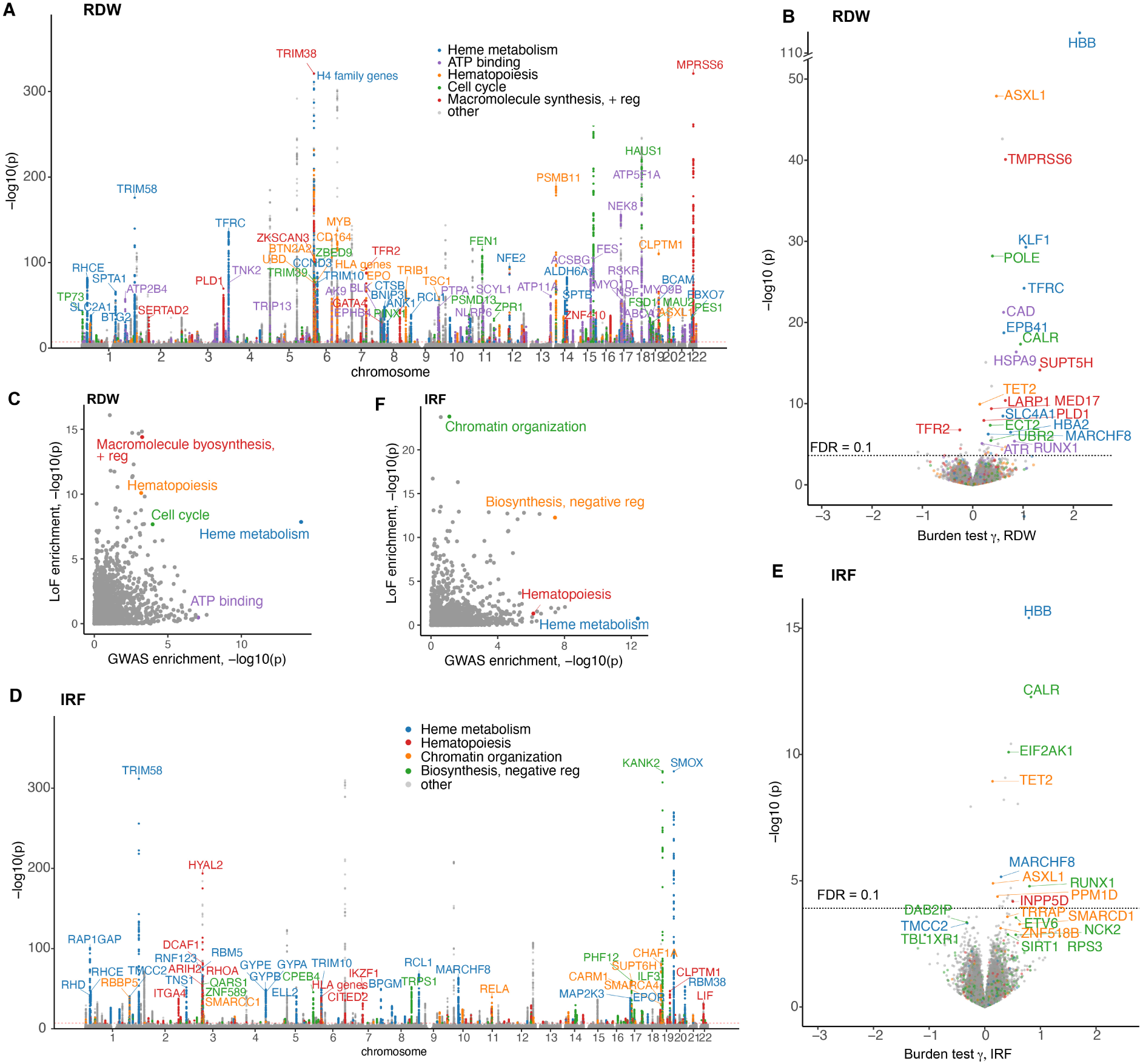
Pathway enrichments for blood trait associations., related to. **Figure 2 A)** Genetic associations identified from UKB GWAS for RDW. Variants located within a 100 kbp window centered on the transcription start site of the genes in the gene set are colored. **B)** Gene association with RDW from UKB LoF burden test. Colors indicate the same gene sets as A). Genes labeled have FDR < 0.01 and belong to the gene sets. **C)** Pathway enrichment of GWAS and LoF burden test top genes for RDW. For GWAS, the closest genes from the independent top variants were used. For the LoF burden test, genes were ranked by the absolute posterior effect size after GeneBayes, and the same number of genes as in GWAS was used. **D)** Genetic associations identified from UKB GWAS for IRF. Variants located within a 100 kbp window centered on the transcription start site of the genes in the gene set are colored. **E)** Gene association with IRF from UKB LoF burden test. Colors indicate the same gene sets as D). Genes labeled have FDR < 0.5 and belong to the gene sets. **F)** Pathway enrichment of GWAS and LoF burden test top genes for IRF. For GWAS, the closest genes from the independent top variants were used. For the LoF burden test, genes were ranked by the absolute posterior effect size after GeneBayes, and the same number of genes as in GWAS was used.

**Figure S3:**
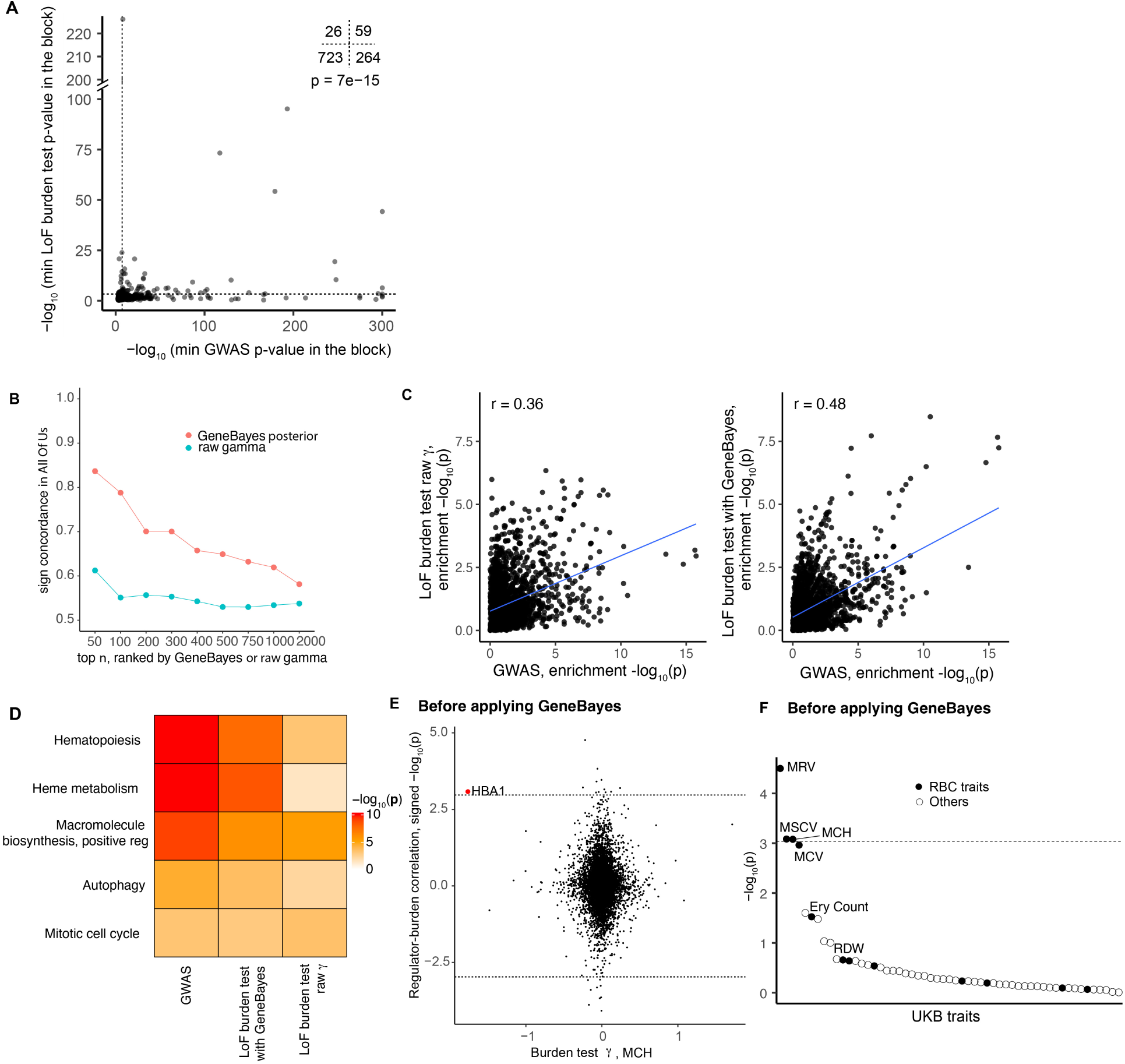
Evaluation of GeneBayes, related to. **Figure 2 and 3. A)** Comparison of GWAS and LoF burden test associations for MCH. We took the minimum GWAS p-value within an LD block, and the minimum LoF burden test p-value for any gene that overlaps the LD block. Dotted lines indicate p = 5 × 10^−8^ *for GWAS and p =* 5 × 10^−4^*, which corresponds to an FDR of 0.1 for the LoF burden test. Numbers of blocks in each quadrant are depicted on the top right corner. p-value is from a two-sided Fisher’s exact test. **B)** Sign concordance of burden test top hits in All Of Us. The result is for RDW. We ranked the genes based on absolute burden test effect size in UKB, either before or after GeneBayes, and assessed the fraction of genes that had the same sign of associations in All Of Us. (C) Enrichment of GO and MsigDb hallmark pathways for genetic associations in MCH, before (left) or after (right) GeneBayes. For GWAS, the closest genes to the lead hits were ordered by p-values. For the LoF burden test, whether or not GeneBayes was applied, genes were ordered by absolute effect sizes. For both GWAS and LoF, the top 200 genes were used for the enrichment analyses. **D)** Enrichment of top 200 genes from GWAS or LoF top hits with or without applying GeneBayes to representative pathways. **E)** Regulator-burden correlation for MCH is compared with their γ for MCH. Same comparison with Figure 3C, but this time using γ before applying GeneBayes. Dotted lines indicate the same threshold with Figure 3C. **F)** Correlation significance of HBA1 regulatory effects with gene effects across a variety of traits. Same comparison with Figure S4A, but this time using γ before applying GeneBayes. Dotted line*

**Figure S4:**
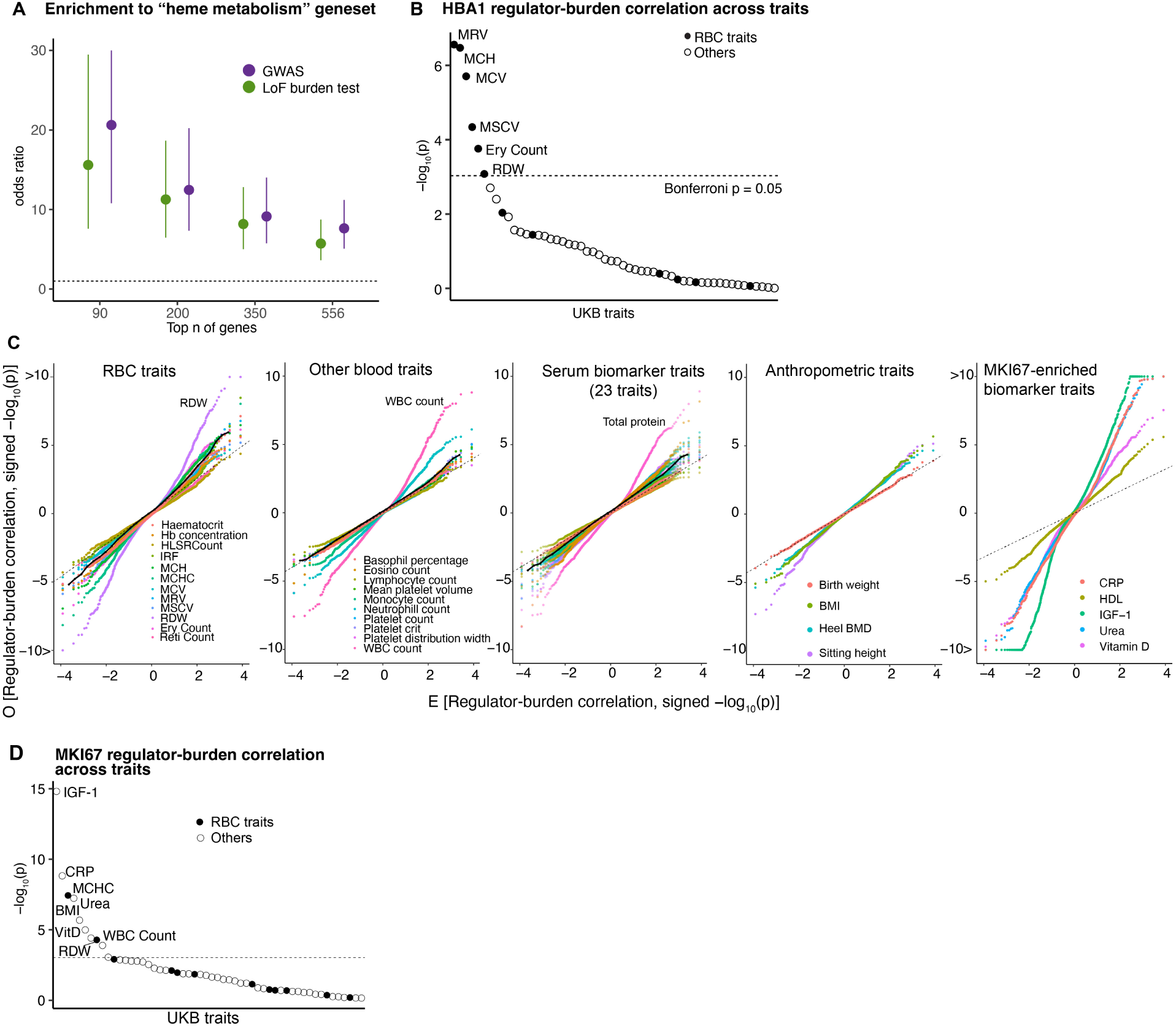
Relevance of gene regulatory effects on trait associations, related to. **Figure 3. A)** Enrichment of hemoglobin metabolism geneset for GWAS and LoF lead hits. For both GWAS closest genes and the LoF burden test, genes were ranked by association p-values, and top gene enrichment for the geneset was assessed using Fisher’s exact test. Error bars indicate 95% confidence intervals. **B)** Correlation significance of HBA1 regulatory effects with gene effects (γ) across a variety of traits. **C)** Genome-wide QQ-plots for burden-regulator correlations for a wide variety of traits. Each dot indicates one gene. Black solid line indicates the median across each category of traits. For serum biomarker traits, 5 traits which showed extensive association with MKI67 regulatory effects are plotted separately. **D)** Correlation significance of MKI67 regulatory effects with gene effects across a variety of traits. Dotted line indicates the threshold for Bonferroni significance.

**Figure S5:**
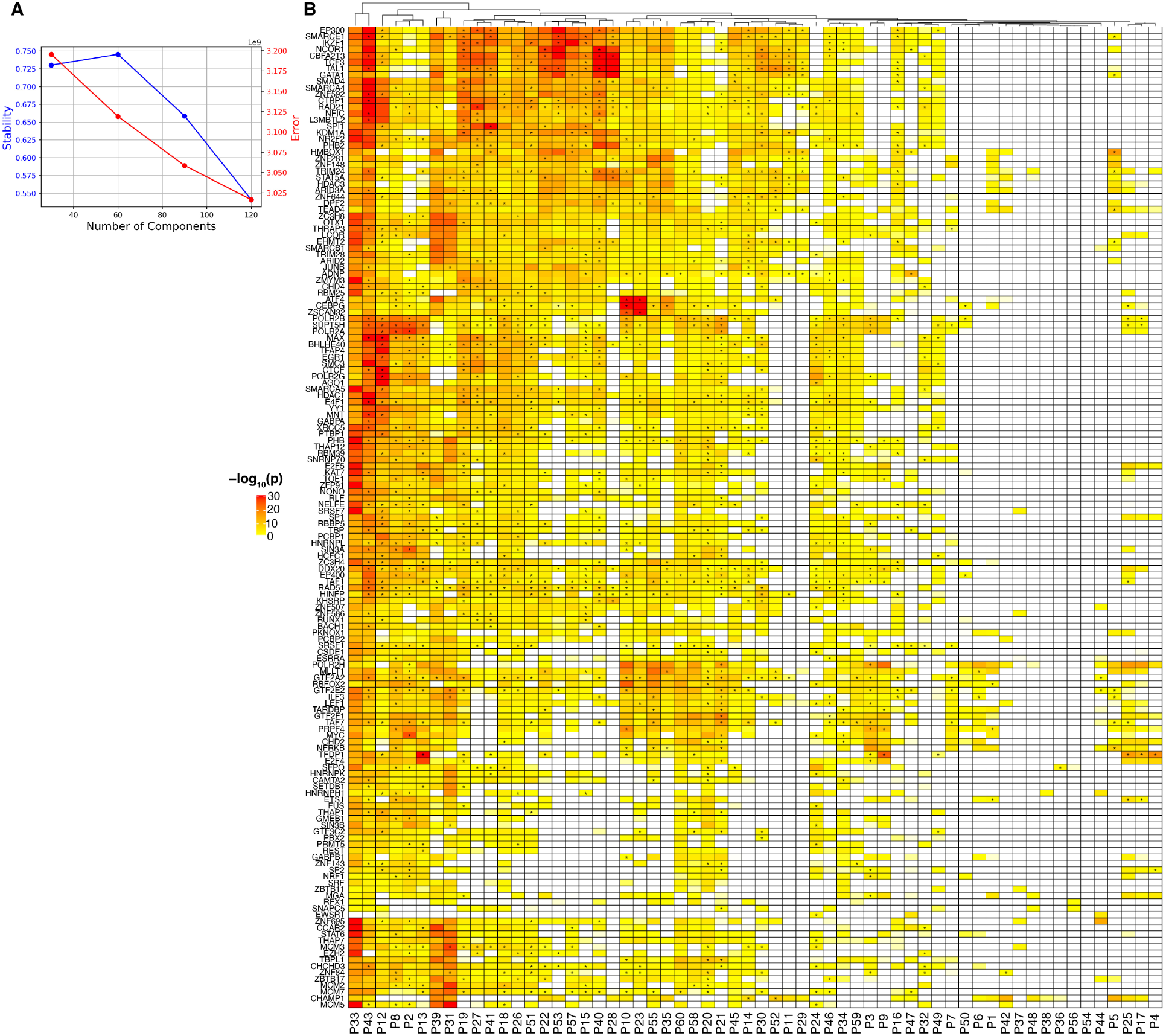
Annotation of programs by transcription factor binding sites, related to. **Figure 4. A)** Number of cNMF components against solution stability measured by the euclidean distance silhouette score of the clustering, and Frobenius error of the consensus solution, outputted by cNMF. **B)** Enrichment of transcription factor binding sites to program genes. Narrow peaks from ChIP-seq of transcription factors (TF) in K562 cells were used to calculate the enrichment (Methods). For significantly enriched TF-program pairs (FDR < 0.05), we tested the effect of knockdown of the TF on program activity and marked an asterisk if the KD also had an effect in the expected direction; that is, if the KD of an activator transcription factor decreased the program activity (p < 0.05), we marked it, and vice versa for repressor.

**Figure S6:**
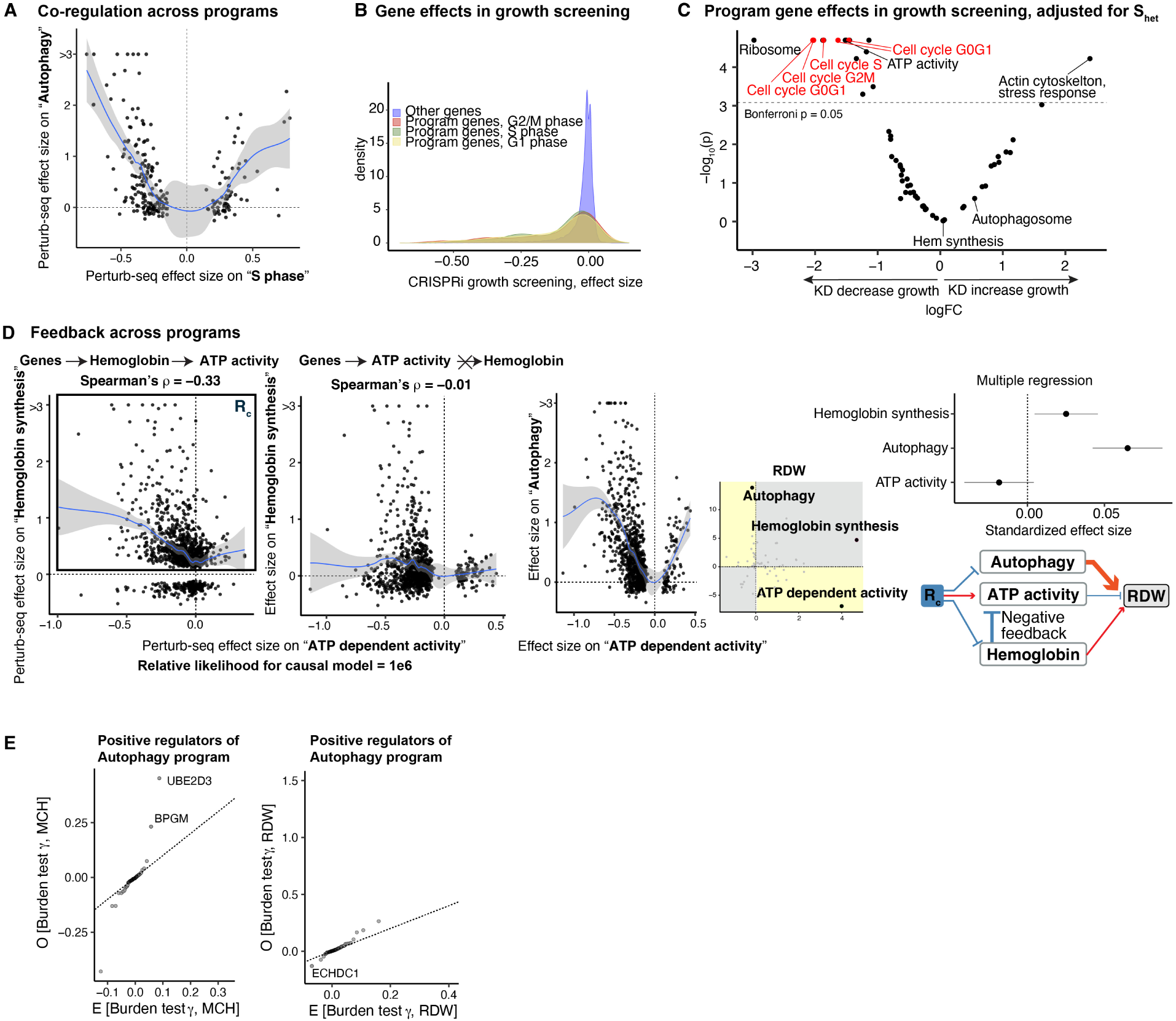
Association of programs and regulators with traits, related to. **Figure 5. A)** Co-regulation pattern between S phase and autophagy programs. Each dot is a gene that has significant regulatory effects on S phase program. **B)** Effects of cell cycle program genes KD on cellular growth. Growth screening data were obtained from an independent experiment using K562 [57]. The effect size is a normalized measure of the impact of KD on cellular growth compared to wild type, denoted as gamma in the original manuscript. **C)** Effects of program genes KD on cellular growth [57]. Here, for each program, we created 100,000 sets of control genes matched for S_het_ and compared the mean effects on cellular growth. **D)** Co-regulation pattern between ATP dependent activity, hemoglobin synthesis and autophagy programs. Genes with regulatory effects on hemoglobin program activity also had effects on ATP activity, but the opposite was not true. Right: Schematics for regulator association with the programs and trait. **E)** Distribution of burden test effect sizes for MCH (left) or RDW (right) for positive regulators of autophagy program. Note that the number of positive regulators is much smaller than that of negative regulators (Figure 5J).

**Figure S7:**
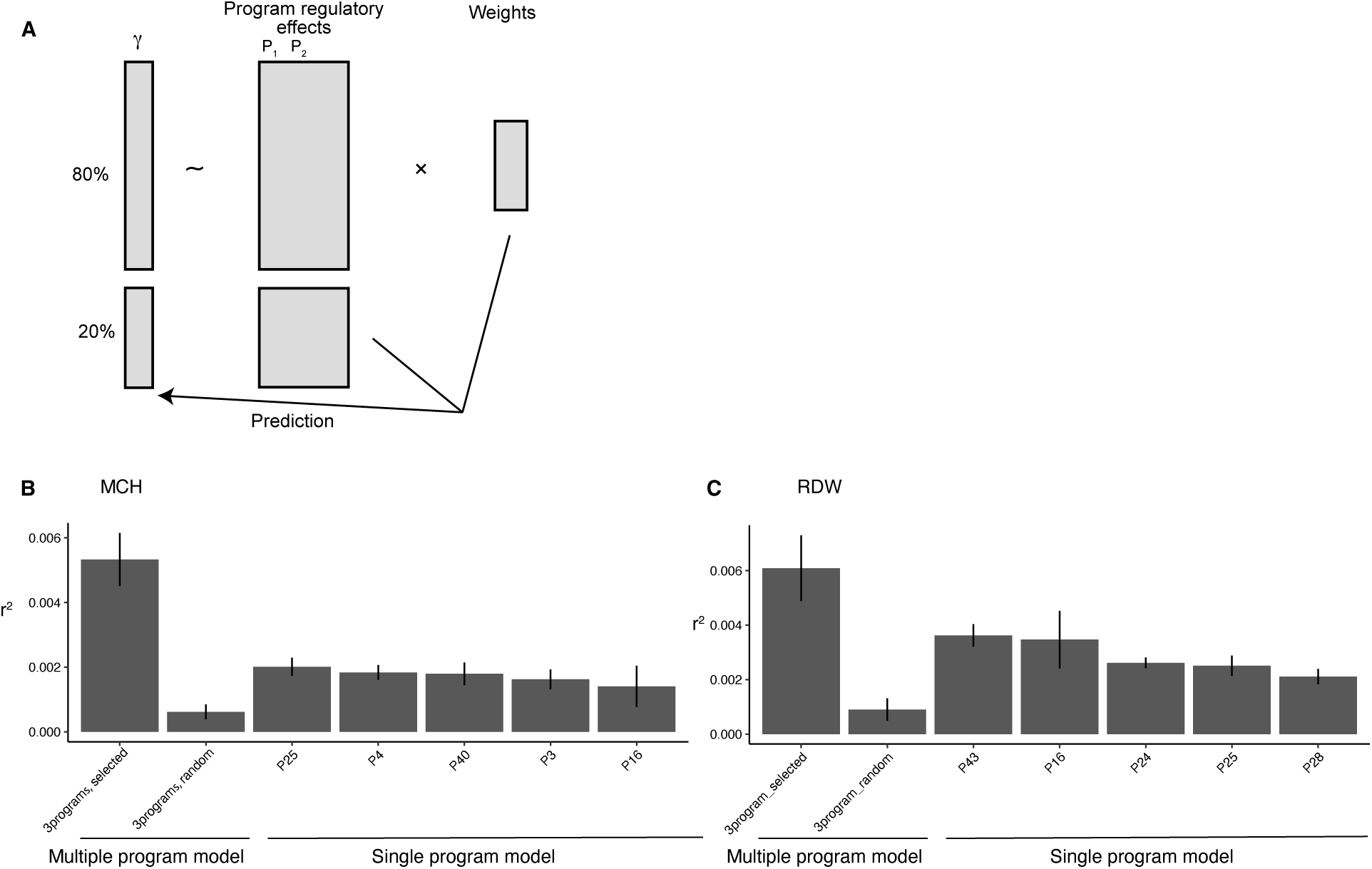
Variance explained by multiple program regulators. **A)** We split the genes into a training set and a test set, and fitted multiple or single regression models to test the association between the gene regulatory effects on the program(s) and the gene effects on the trait (γ). We evaluated the variance explained by the model using the test gene set. **B-C)** Variance explained by the regression models for MCH (B) and RDW (C). “Programs, selected” refers to the programs selected from the regulator-burden correlations in the gene-to-program-to-trait map. “Programs, random” refers to the randomly selected sets of multiple programs. Single programs shown are the top 5 programs as to the variance explained. Error bars indicate 1.96× *standard errors*.

**Figure S8:**
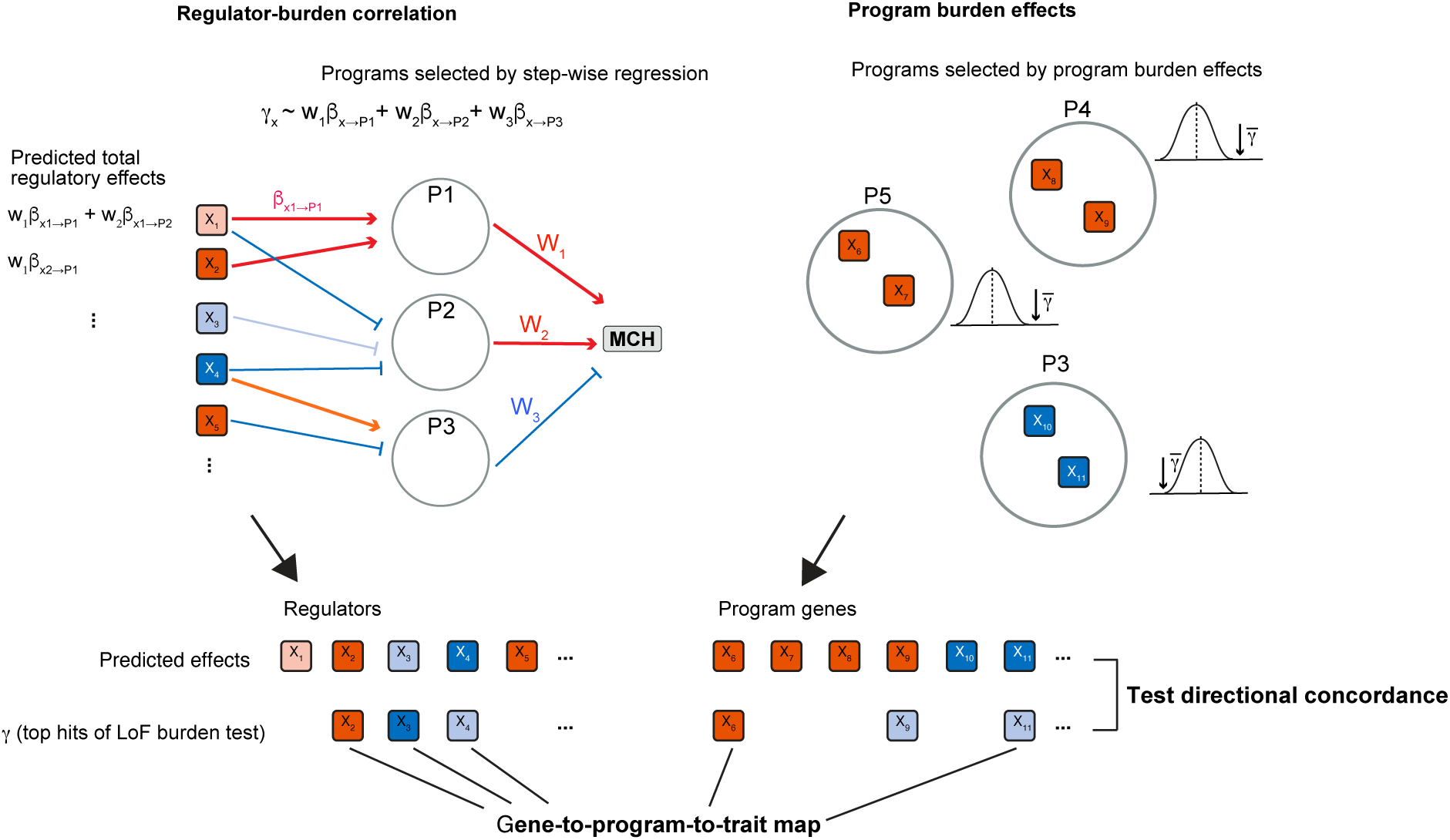
Schematics for making multiple program association model.

**Figure S9:**
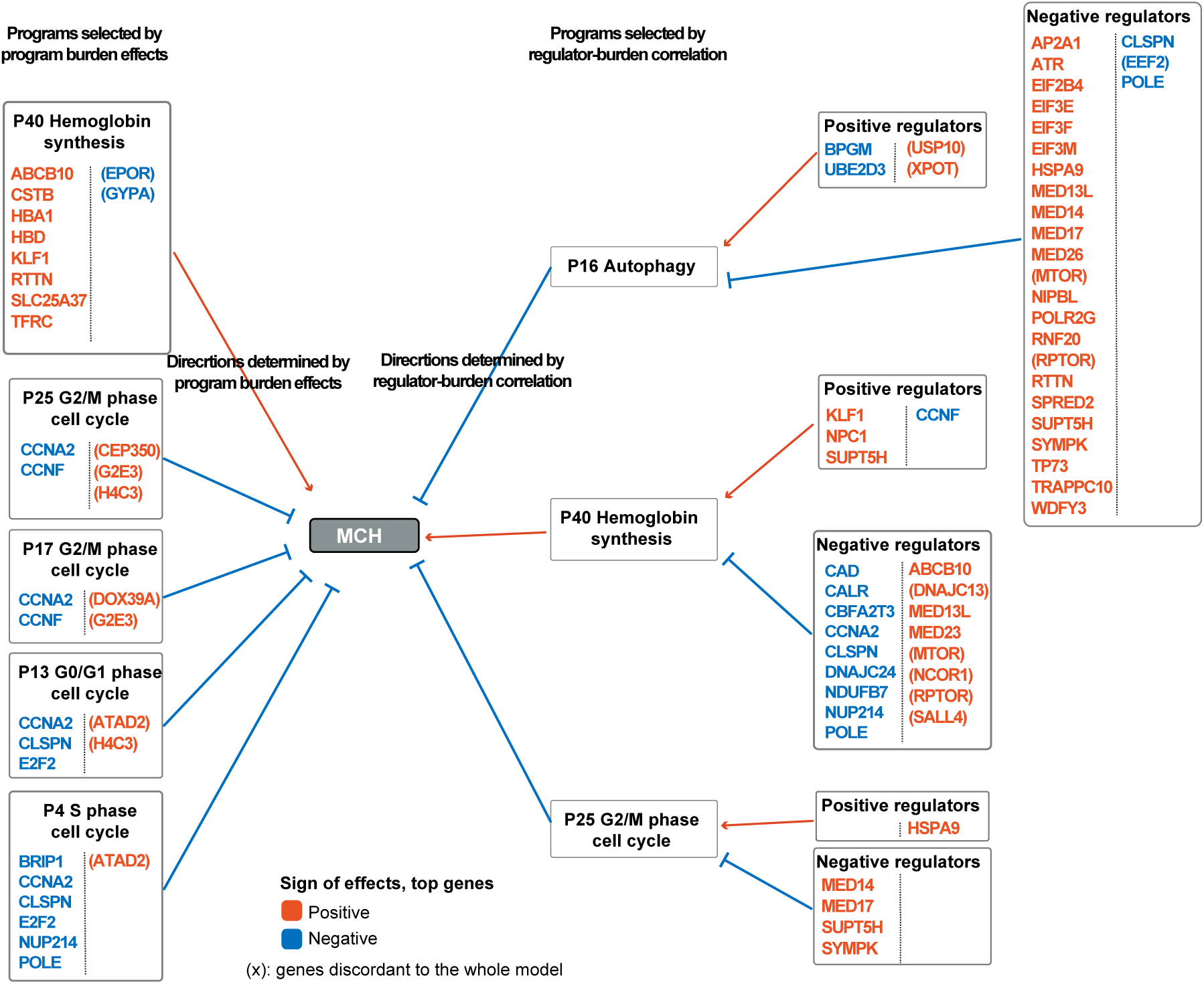
Programs selected for modeling MCH associations. Programs were selected based on program burden effects (left) or regulator-burden correlations (right). For each program, top hits (|*γ*| > 0.1*) for MCH that overlap with the top 200 loading genes (for program genes) or regulators (FDR < 0.05, for regulator genes) are listed. The color of genes correspond to the sign of γ. Genes in parentheses are discordant from the predicted directions from the overall model. Some of the genes are associated with multiple programs or regulators*.

**Figure S10:**
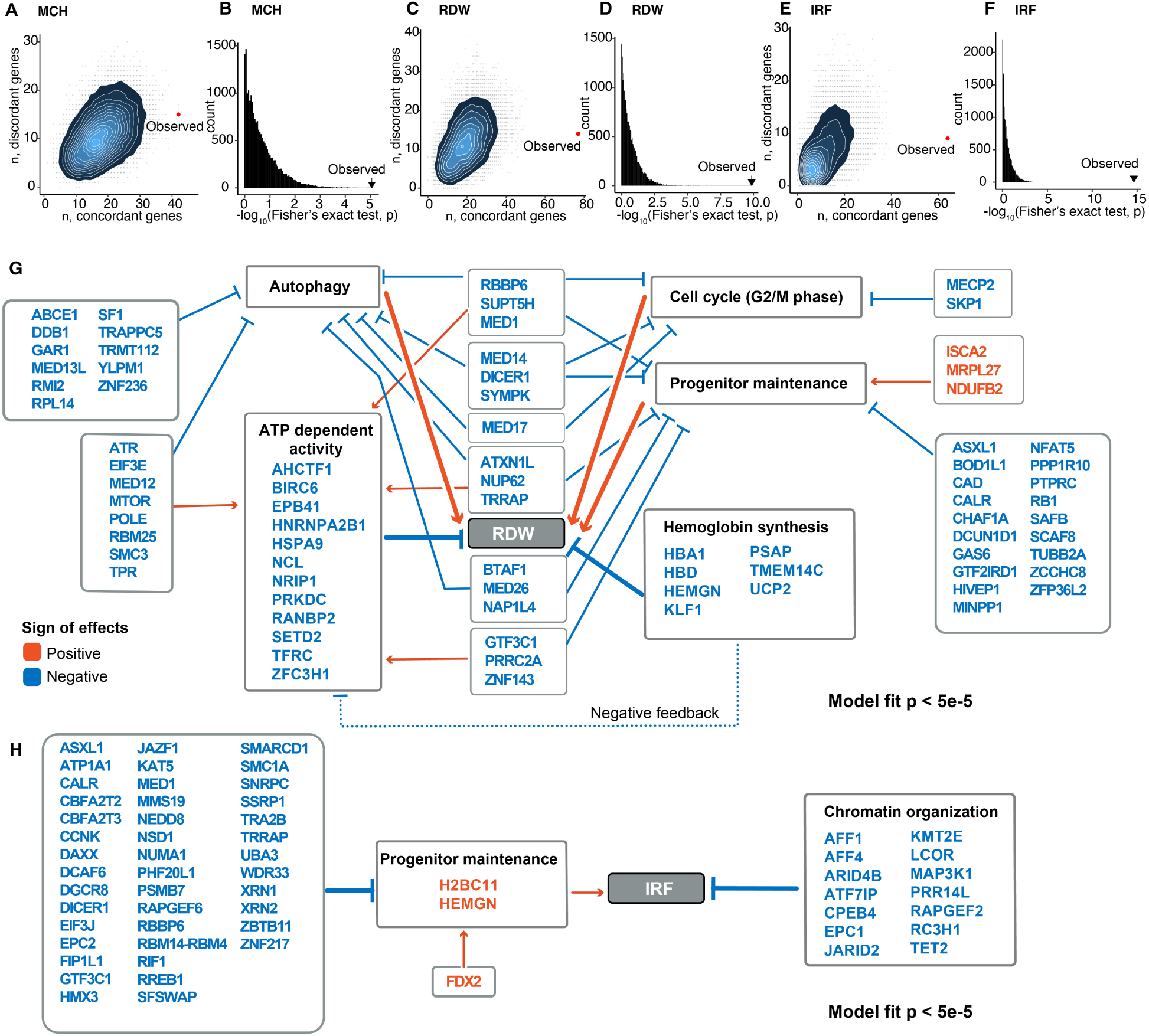
Gene to program to trait maps, related to. **Figure 6. A)** Number of top hits (|*γ*| > 0.1*) for MCH whose direction of associations were concordant or discordant with that predicted from the model. Grey points and their density plot are the results from 20,000 permutations. Red point shows the observed data. **B)** Distribution of top hits concordance p-values in permutation tests for MCH. In each permutation, we counted the number of top hits concordant with the model and evaluated its enrichment (Methods). The observed result showed the highest concordance compared to permuted sets. **C-F)** Same plots as (A) and (B), for RDW (C,D) and IRF (E, F). **G-H)** Gene to program to trait map for RDW (G) and IRF (H)*.

**Figure S11:**
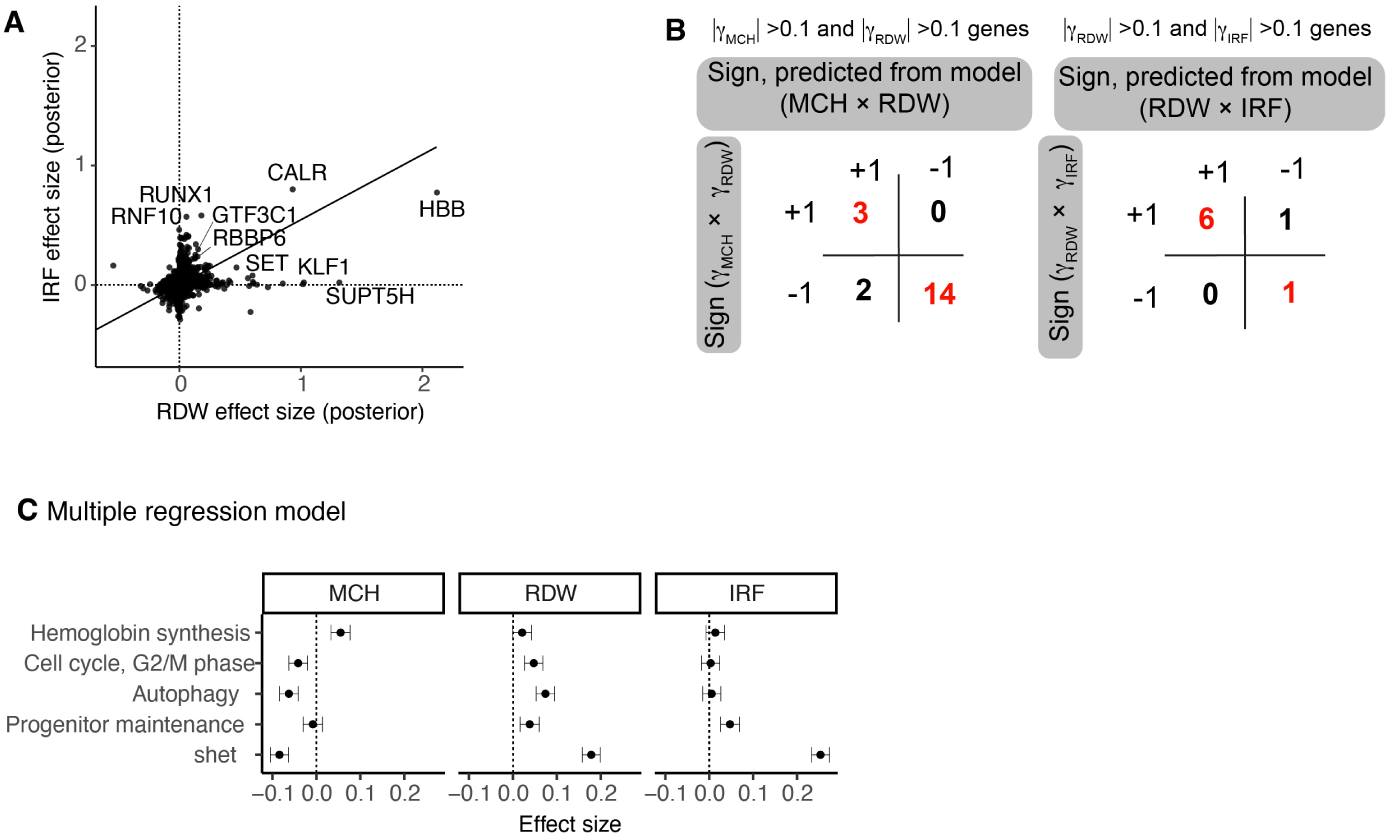
Cross-trait comparisons of gene effects, related to. **Figure 6. A)** Comparison of LoF burden test effect sizes after GeneBayes between IRF and RDW. The solid line corresponds to the first principal component. **B)** Cross-trait directional relationships of gene effects in the predicted gene-to-program-to-trait model and raw data from the LoF burden test. The left table shows the comparison between MCH and RDW, while the right table shows the comparison between RDW and IRF. For each table, only genes that have strong effects in both traits (|*γ*| > 0.1*) and selected in the predicted model for both traits are considered. For instance, +1 means that the gene has strong effects for both traits in the same direction. MCH and IRF share few genes with strong effects and could not be compared. **C)** Correlation of regulatory effects on four programs or shet with γ. For each trait, correlation coefficients were estimated with the multiple regression model. Error bars indicate* 95% *CI*.

**Figure S12:**
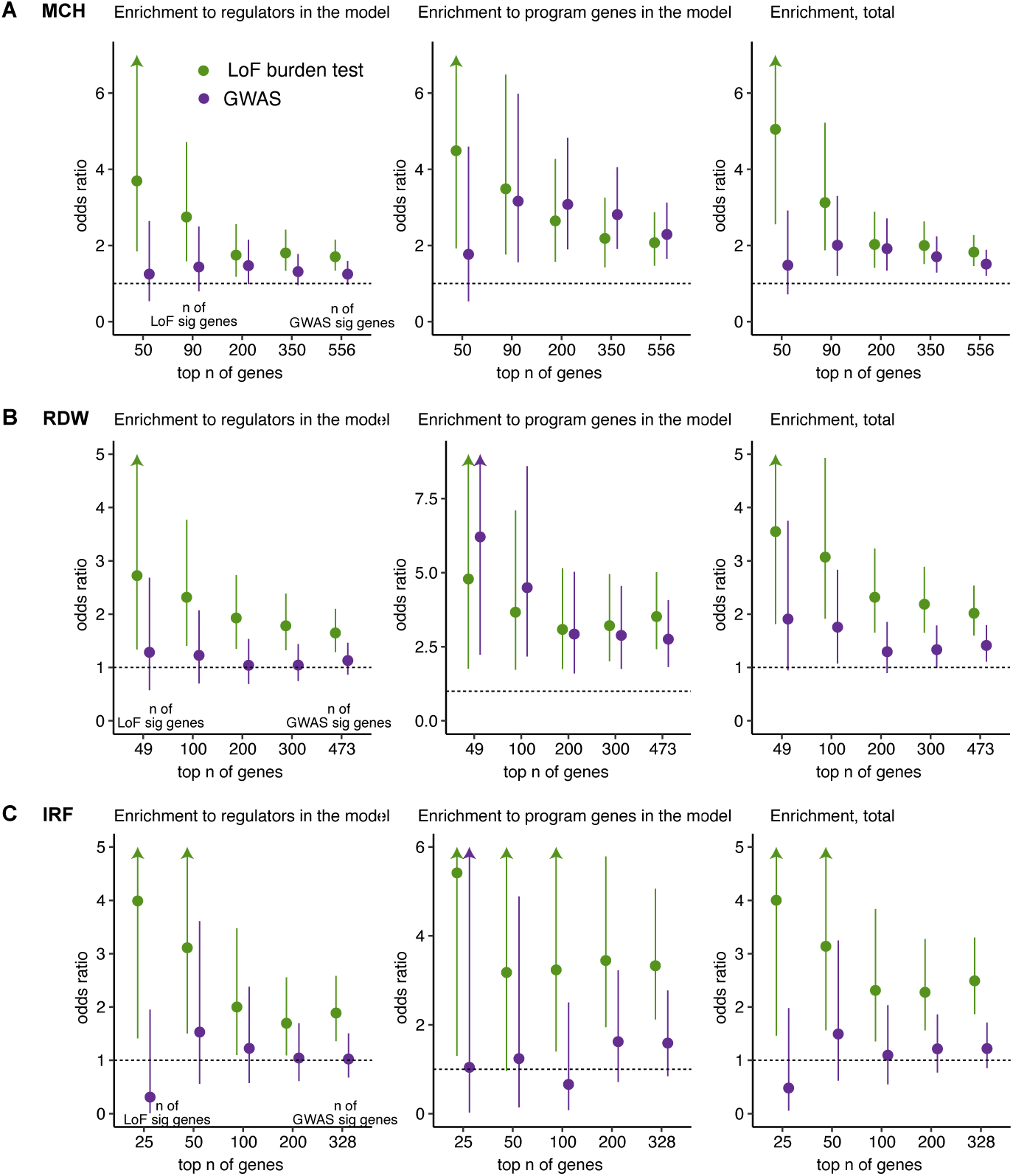
Enrichment of GWAS and LoF top hits in the regulators and program genes in the model. A-C) From the gene-to-program-to-trait map for MCH (A), RDW (B) or IRF (C), we divided the genes into regulators, which have significant regulatory effects on the selected program, and program genes, which are among the top 200 loadings of the selected programs. We ranked the genes closest to the GWAS top hits based on p-values and the genes with LoF burden test based on absolute γ. Using different thresholds for top genes, we tested the enrichment in the genes selected by the model using Fisher’s exact test. “Total” refers to the results for program and regulator genes combined.

**Figure S13:**
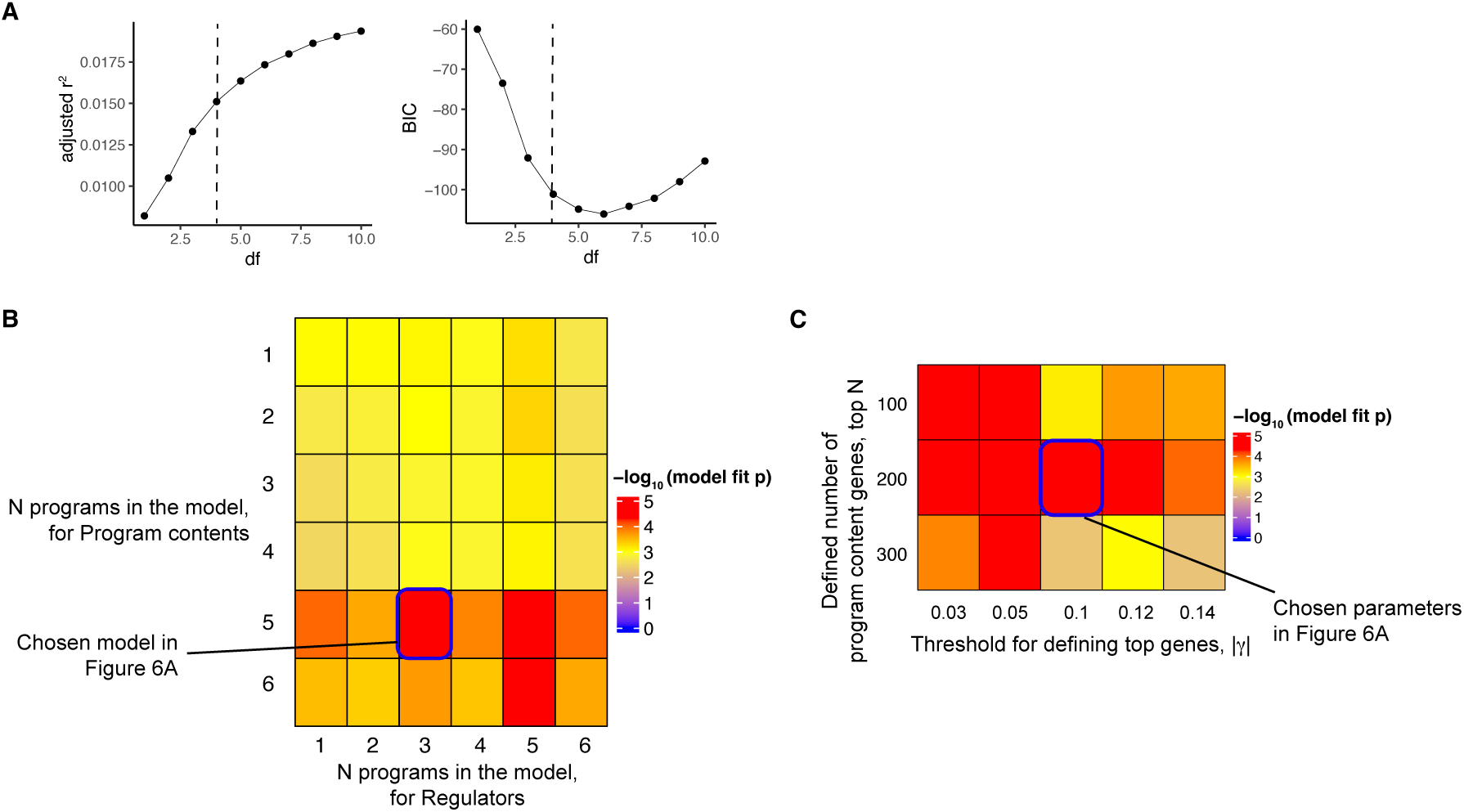
Choice of parameters for gene-to-program-to-trait map. **A)** Adjusted r^2^ and Bayesian Information Criterion (BIC) in the step-wise joint regression model. This is a result for selecting the number of programs whose regulators are associated with MCH. Dotted lines indicate the number of parameters chosen in the model. **B)** Model fit p-value, estimated from 20,000 permutations, for different numbers of programs included in the model for MCH. We varied the number of programs selected from regulator-burden correlations (x-axis) and program burden effects (y-axis) and evaluated the fit of the top hits to the model. **C)** Model fit p-value for different definitions of program genes or top hit genes from the permutation test. Here, we tested for MCH with the same number of programs in the final model.

